# Data-driven characterization of Preterm Birth through intramodal Diffusion MRI

**DOI:** 10.1101/2023.01.12.523771

**Authors:** Rosella Trò, Monica Roascio, Domenico Tortora, Mariasavina Severino, Andrea Rossi, Eleftherios Garyfallidis, Gabriele Arnulfo, Marco Massimo Fato, Shreyas Fadnavis

## Abstract

Preterm birth still represents a concrete emergency to be managed and addressed globally. Since cerebral white matter injury is the major form of brain impairment in survivors of premature birth, the identification of reliable, non-invasive markers of altered white matter development is of utmost importance in diagnostics. Diffusion MRI has recently emerged as a valuable tool to investigate these kinds of alterations. In this work, rather than focusing on a single MRI modality, we worked on a compound of beyond-DTI High Angular Resolution Diffusion Imaging (HARDI) techniques in a group of 46 preterm babies studied on a 3T scanner at term equivalent age and in 23 control neonates born at term. After extracting relevant derived parameters, we examined discriminative patterns of preterm birth through (i) a traditional voxel-wise statistical method such as the Tract-Based Spatial Statistics approach (TBSS); (ii) an advanced Machine Learning approach such as the Support Vector Machine (SVM) classification; (iii) establishing the degree of association between the two methods in voting white matter most discriminating areas. Finally, we applied a multi-set Canonical Correlation Analysis (CCA) in search for sources of linked alterations across modalities. TBSS analysis showed significant differences between preterm and term cohorts in several white matter areas for multiple HARDI features. SVM classification performed on skeletonized HARDI measures produced satisfactory accuracy rates, especially as for highly informative parameters about fibers’ directionality. Assessment of the degree of overlap between the relevant measures identified by the two methods exhibited a good, though parameter-dependent rate of agreement. Finally, CCA analysis identified joint changes precisely for those features exhibiting less correspondence between TBSS and SVM. Our results suggest that a data-driven intramodal imaging approach is crucial to extract deep and complementary information that cannot be extracted from a single modality.

## INTRODUCTION

Despite the development of monitoring and treatment technology in the neonatal intensive care unit, the incidence of preterm birth is still increasing worldwide (Beck et al., 2010; Blencowe et al., 2013), with a high rate of neurodevelopmental impairment among preterm survivors (Bhutta et al., 2002). These sequelae have been attributed to perinatal brain injury, with the severity of White Matter (WM) injury being predictive of neurodevelopmental outcome in later childhood (Kimpton et al., 2021; Dyet et al., 2006; Tortora et al., 2018). Thus, increasing the understanding of WM microstructural development between birth and Term Equivalent Age (TEA) in preterm infants might be informative of whole-brain maturational processes in early life, both in case of normal or adverse neurodevelopmental outcomes.

As such cerebral alterations often occur beneath an anatomical scale, being therefore undetectable by conventional imagery, diffusion MRI (dMRI) has been established as the most valuable tool to inspect this altered WM development in preterm infants (Volpe, 2003; Counsell et al., 2003). For instance, previous neonatal brain Diffusion Tensor Imaging (DTI) studies indicated that WM Fractional Anisotropy (FA) increases with age, even before myelin is evident on conventional MR imaging sequences, and diffusion parameters correlate with cognitive, language, and motor outcomes (Hüppi et al., 1998a; Partridge et al., 2004). If examination of WM development in preterm neonates by using DTI is by then recognized as a trusted tool to identify neonates at high risk of neurodevelopmental impairment, the recent advent of High Angular Resolution Diffusion Imaging (HARDI) represents a promising implementation for enhancing our understanding of WM maturation compared with standard DTI metrics alone (Kunz et al., 2014). Most recent studies show how higher-order diffusion models (e.g., Diffusion Kurtosis Imaging (DKI) (Zhao et al., 2021; Ouyang et al., 2019b; Shi et al., 2016), Neurite Orientation Dispersion and Density Imaging (NODDI) (Kelly et al., 2016) or Constrained Spherical Deconvolution (CSD) (Pannek et al., 2018)) may offer unique insight into the postnatal neurological development associated with differential degrees of preterm birth, thanks to their improved sensitivity to microstructural changes.

Indeed, by relating the diffusion signal more directly and specifically to underlying cellular microstructural properties, these HARDI methods allow to investigate WM microstructure with higher levels of detail and to potentially give access to richer microstructural properties, thus increasing the diagnostic values of their derived measures. Cutting-edge approaches of microstructural brain imaging are thus expected to provide unprecedented insights into early brain anomaly leading to subsequent adverse outcomes, often intrinsic to very preterm birth and whose accurate prediction is imperative for the implementation of early interventions in clinical practice.

However, as far as we are aware, all studies using HARDI techniques investigate preterm-related brain injuries resorting to one microstructural model at a time. Conversely, an overview of multiple models may be extremely useful to offer complementary information about tissue microstructure, given also the lack of systematic model-to-model comparisons in search of the best markers to reliably quantify microstructural tissue maturation in newborns. Specifically, in this study, we considered four models among the most used in paediatrics: DKI (Jensen et al., 2005), NODDI (Zhang et al., 2012), Multi-Shell Multi-Tissue CSD (MSMT CSD) (Jeurissen et al., 2014) and Fiber Orientation Estimated using Continuous Axially Symmetric Tensors (FORECAST) (Anderson, 2005; Kaden et al., 2016) selected for their suitability in grasping microstructural changes beyond DTI’s capabilities (Pecheva et al., 2018).

Our work thus primarily aims at investigating the potential of an intramodal approach to dMRI, made up of several microstructural dMRI models, in extracting relevant markers of prematurity. Furthermore, with a view to deeply inspecting each of the microstructural-derived measures, we also seek to assess their robustness to the method of investigation in use. To do so, we opted for comparing two state-of-the-art methods for large-scale longitudinal neuroimaging studies, based on Tract-Based Spatial Statistics (TBSS) and Machine Learning (ML) classification, respectively.

TBSS is an automated, observer-independent approach for assessing FA in the major WM tracts on a voxel-wise basis across groups of subjects (Smith et al., 2006). It achieves this through carefully tuned alignment of FA maps to a standard-space template, followed by projection of individual data onto a skeletonised representation of major WM tracts common to the group to circumvent the Partial Volume Effect (PVE) and gain statistical power from this dimensionality reduction. Since WM dysmaturation is increasingly recognized as the primary pathology in contemporary cohorts of preterm neonates, this method has been used extensively on scans acquired at TEA in preterm-born neonates to successfully detect alterations in WM microstructure in the absence of overt brain injury (Anjari et al., 2007; Alexandrou et al., 2014) and to predict cognitive and motor outcomes in young preterm-born children (Tortora et al., 2018; Duerden et al., 2015; Counsell et al., 2002), which is highly relevant to clinicians making essential care decisions. The reasons behind the popularity of TBSS reside in being an objective, sensitive and relatively easy-to-interpret method for multi-subject, whole-brain diffusion data analysis. This allows overcoming some of the limitations of common ROI-based approach to analyze neonatal data, suffering from subjectivity, manual intensity, intra- and inter-subject variability, and a priori spatial localization, which makes it suboptimal for comparison of several brain regions or large subject groups (Ly et al., 2015). At the same time, the approach could alleviate many concerns raised regarding other conventional fully automated whole-brain measurement techniques such as the Voxel-Based Morphometry (VBM) framework, previously used in many DTI studies (Whitwell, 2009). Although remaining the leading technique for voxel-wise DTI analysis, as with any other voxel-based analysis method, the application of TBSS does not come without pitfalls, including the influence of noise level, parameter settings, choice of the template, quality of image registration on the resulting anatomical specificity, low sensitivity for detecting wide-spread subtle abnormalities, impossibility to develop individually-based imaging indices. All these factors in turn question the reproducibility and robustness of the final TBSS result, being essential for establishing biomarkers and diagnostic/prognostic indices at the individual level (Bach et al., 2014).

In this respect, the advent of ML in the early 2000s has totally revolutionized neuroimaging studies. Indeed, compared to population-based analysis, ML approaches can extract unbiased, individualized biomarkers of diseases or functional brain states of fundamental importance in diagnosis, prognosis, and patient stratification (Davatzikos, 2019). Earlier studies have focused on Support Vector Machines (SVM) (Golland et al., 2002; Lao et al., 2004), which has been a cornerstone in this field, largely because of its robustness and ease of use with a variety of kernels (Schölkopf et al., 2002). These and other methods have already been widely applied in neuroimaging studies regarding preterm birth (Chu et al., 2015; Galdi et al., 2020; Saha et al., 2020), with the vast majority of works resorting to TBSS or other VBM methods as a preprocessing step preparatory to application of ML algorithms. However, to the best of our knowledge, this is the first study using TBSS and an ML-based classification jointly on intramodal dMRI to explore the most discriminating WM regions as biomarkers to supplement the understanding of such a current phenomenon as preterm birth.

Moreover, given the multiplicity of microstructural measures examined, we considered it informative also to apply a Canonical Correlation Analysis (CCA) approach (Hardoon et al., 2004), which focuses on finding linear combinations that account for the most correlation in two or more datasets. This method has proved to outperform traditional statistical tools in unlocking the complex relationships among many variables in large datasets (Wang et al., 2020). Indeed, it has been proven that different dMRI metrics themselves share overlying information potentially causing partial redundancies in data analysis, especially in the case of neonatal imaging (Chamberland et al., 2019; Cox et al., 2016; De Santis et al., 2014; Girault et al., 2019). This in turn hints that the identification of a reduced number of microstructurally informative and biologically-interpretable components may turn out to be particularly useful. Relating to our survey, CCA is specifically beneficial to capitalize on the strength of each microstructural imaging feature so as to uncover hidden relationships among separate microstructural models. Despite its multiple applications in medical imaging (Correa et al., 2008, 2010a,b; Hardoon et al., 2007; Sui et al., 2011, 2013) to the best of our knowledge, no study has used intramodal advanced dMRI data fusion to examine of the full relationships among multiple dMRI models to provide more informative insights into altered brain patterns typical of prematurity. Understanding the joint changes in different advanced dMRI approaches related to premature birth may advance our knowledge of underpinning microstructural mechanisms and provide potential intramodal biomarkers for better clinical diagnosis.

## METHODS AND MATERIALS

### Subjects

A total of forty-six preterm and twenty-three term-born subjects have been enrolled between November 2017 and August 2021 at the Neuroradiology Unit of Gaslini Children’s Hospital. Conventional MRI and DKI were performed using a 3.0T MRI scanner (Ingenia Cx, Philips, Best, the Netherlands) with a 32-channel head array coil.

This single-centre study was carried out in accordance with the recommendations of “Comitato Etico Regione Liguria, Genoa, Italy” with written informed parental consent obtained for each infant prior to examination in accordance with the Declaration of Helsinki. All the precautions for neonatal patient feeding, mild sedation, position and monitoring have been adopted. Exclusion criteria included obvious motion artefacts, oblique positioning, an incomplete imaging process or a low SNR.

GA was used as a classifying variable for preterm (GA < 37 weeks) and term birth (GA ≥ 37 weeks). Preterm subjects have been acquired at TEA, for clinical practice. Preterm and term neonates did not show significant brain abnormalities at visual inspection on T2- and T1-weighted images. Details about subjects’ demographics are reported in Table 1.

**Table 1.**
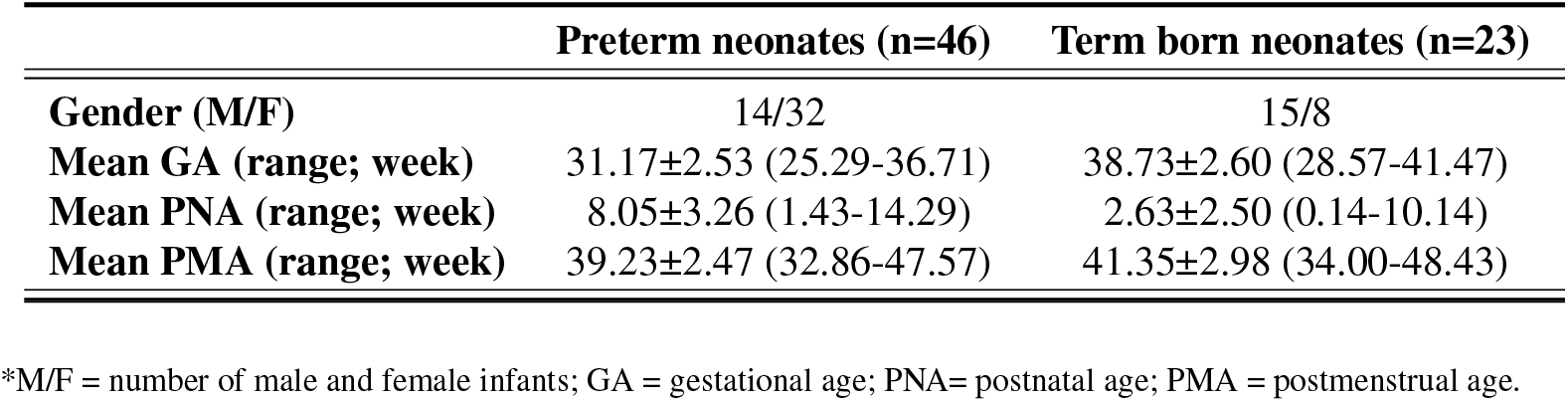
Demographic features of neonates-brain

|  | Preterm neonates (n=46) | Term born neonates (n=23) |
| --- | --- | --- |
| **Gender (M/F)** | 14/32 | 15/8 |
| **Mean GA (range; week)** | 31.17±2.53 (25.29-36.71) | 38.73±2.60 (28.57-41.47) |
| **Mean PNA (range; week)** | 8.05±3.26 (1.43-14.29) | 2.63±2.50 (0.14-10.14) |
| **Mean PMA (range; week)** | 39.23±2.47 (32.86-47.57) | 41.35±2.98 (34.00-48.43) |

\*M/F = number of male and female infants; GA = gestational age; PNA= postnatal age; PMA = postmenstrual age.

#### MR Acquisition

Our acquisition protocol includes Turbo Field Echo (TFE) 3D T1-weighted and HARDI sequences. Details about acquisition are reported in Table 2.

**Table 2.** Acquisition protocols for structural T1 and HARDI series

|  | 3dT1 | HARDI |
| --- | --- | --- |
| **TR/TE (s)** | 0.6/0.026337 | 2.086/0.114 |
| **Diffusion Scheme (s/mm<sup>2</sup>)** | – | 5 b=0, 30 b=700, 60 b=2800 |
| **Flip Angle (°)** | 90 | 90 |
| **Reconstruction Resolution (mm)** | 0.38*0.38 | 1.5*1.5 |
| **Reconstruction Matrix** | 512*512 | 144*144 |
| **Multi-Band Factor** | – | 2 |
| **# Averages** | 2 | 1 |
| **Slice Thickness (mm)** | 0.5 without gap | 2.2, without gap |
| **Slice Orientation** | sagittal | axial |
| **# Slices** | 251 | 42 |
| **Total Scan Time** | 4 minutes 5 s | 3 minutes 30 s |
| **Partial Fourier factor** | – | 0.6 |

#### Preprocessing pipeline

##### Structural Images

The first critical step was skull-stripping. Dealing with neonatal scans, standard skull-stripping methods (Hosseini et al., 2015;Smith, 2000;Iglesias et al., 2011;Shattuck and Leahy, 2000) failed in correctly removing non-brain areas, thus requiring manual corrections and introducing both a user- and a subject-based bias. Therefore, we resorted to MASS (Multi Atlas Skull Stripping) (Doshi et al., 2013), which performs brain extraction through a template selection strategy obtaining higher accuracy than recent state-of-the-art tools and avoiding user’s intervention. As a preliminary step, 3D T1-weighted images were FOV-reduced, processed with Brain Extraction Toolbox (BET), and then bias-field corrected with N4 algorithm to suppress low-frequency inhomogeneities (Tustison et al., 2010). At this phase, under supervision of a board-certified neuroradiologist, we selected six subjects that best represented the anatomical variations within the dataset, and processed this cohort with the developing Human Connectome Project (dHCP) pipeline (Hughes et al., 2017). The six 3D T1-weighted betted images generated with dHCP pipeline were subsequently used as a reference template to train the MASS algorithm. A final re-run of the N4 algorithm ensured bias-field correction using the correct mask extracted with MASS framework instead of the rough one after preliminary brain extraction with BET. All preprocessing relative to structural scans are summarized in Figure 1A.

**Figure 1.**
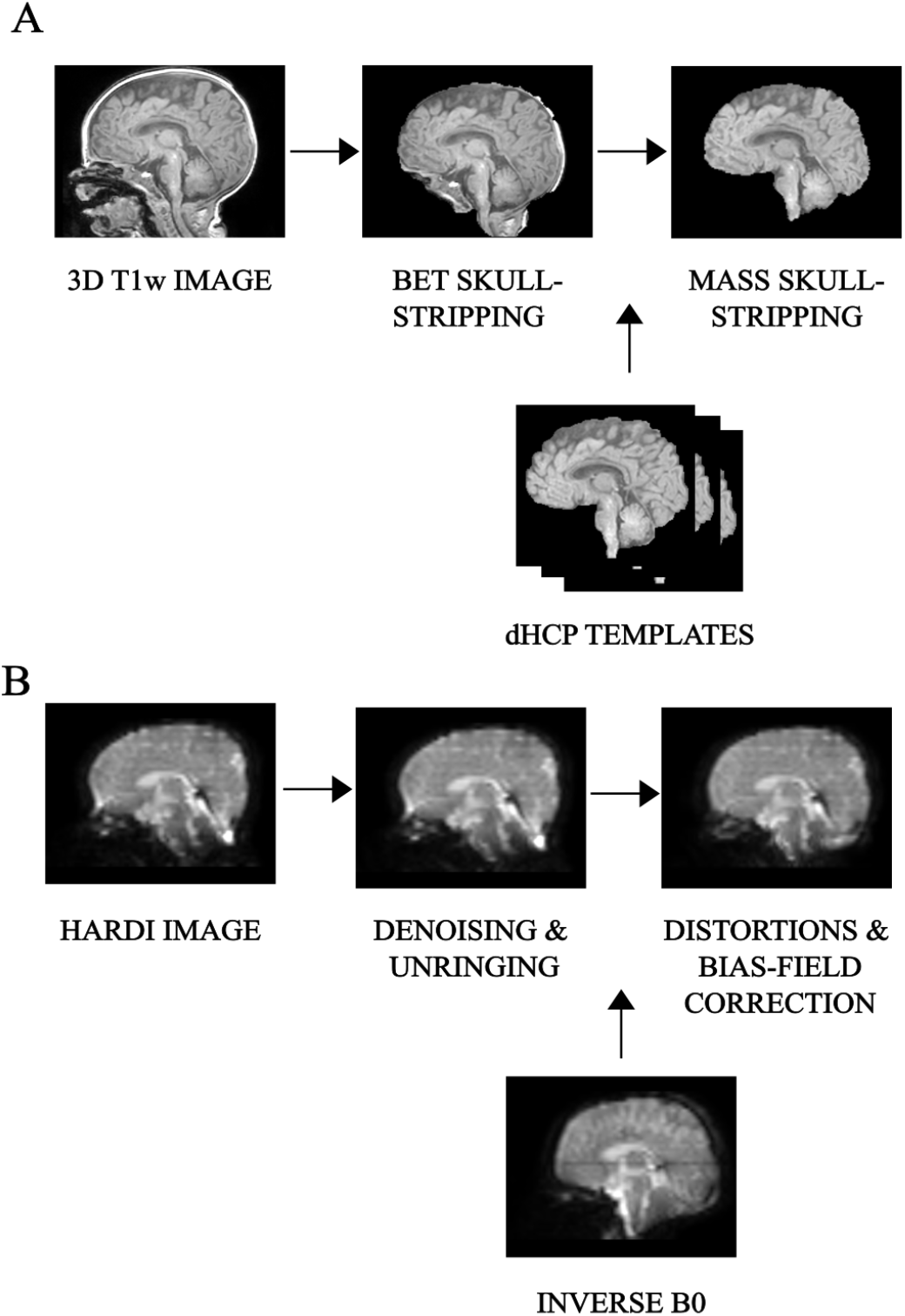
Preprocessing pipeline: overview of the main preliminary image processing steps performed on: **(A)** 3D T1-weighted, whose key step is skull-stripping and **(B)** HARDI scans, whose core is represented by denoising as well as distortion correction, for an example subject.

##### HARDI scans

HARDI scans in paediatrics are really sensitive to low Signal-To-Noise Ratio (SNR), and more prone to macro as well as micro sources of movements. We thus resorted to Patch2Self denoising (Fadnavis et al., 2020) as the very first preprocessing step regarding diffusion imaging. This denoiser turned out to be particularly suitable for higher-order diffusion models, outperforming other existing methods at visual and modelling tasks (Schilling et al., 2022). The method was implemented in DIPY v.1.4.0 (Garyfallidis et al., 2014) and applied with OLS regressor, with the threshold for *b* = 0 shell at 100, given the variability of non-diffusion-weighted *b* values. All subsequent preprocessing steps were done in Mrtrix3 v.3.0.1 (Tournier et al., 2019). Standard analysis pipeline performed well also on neonatal scans thanks to overall good image contrast - (i) denoising; (ii) unringing; (iii) EPI-distortion correction, eddy-current and movement distortion correction; (iv) B1-field inhomogeneity correction. All preprocessing relative to diffusion images is displayed in Figure 1B.

##### Microstructural models

This was the starting point for the application of multiple advanced microstructural dMRI models, easily employed to this cohort thanks to the overall high image quality (Figure 2). The outcome produced by each model has been subject to inspection by two experienced neuroradiologists and compared with existing studies on age-matched cohorts. Furthermore, to avoid spurious contributions from non-representative image portions as well as to reduce computational time, all models have been applied to a masked version of the data derived from averaging and skull-stripping the non-diffusion weighted pre-processed volumes.

**Figure 2.**
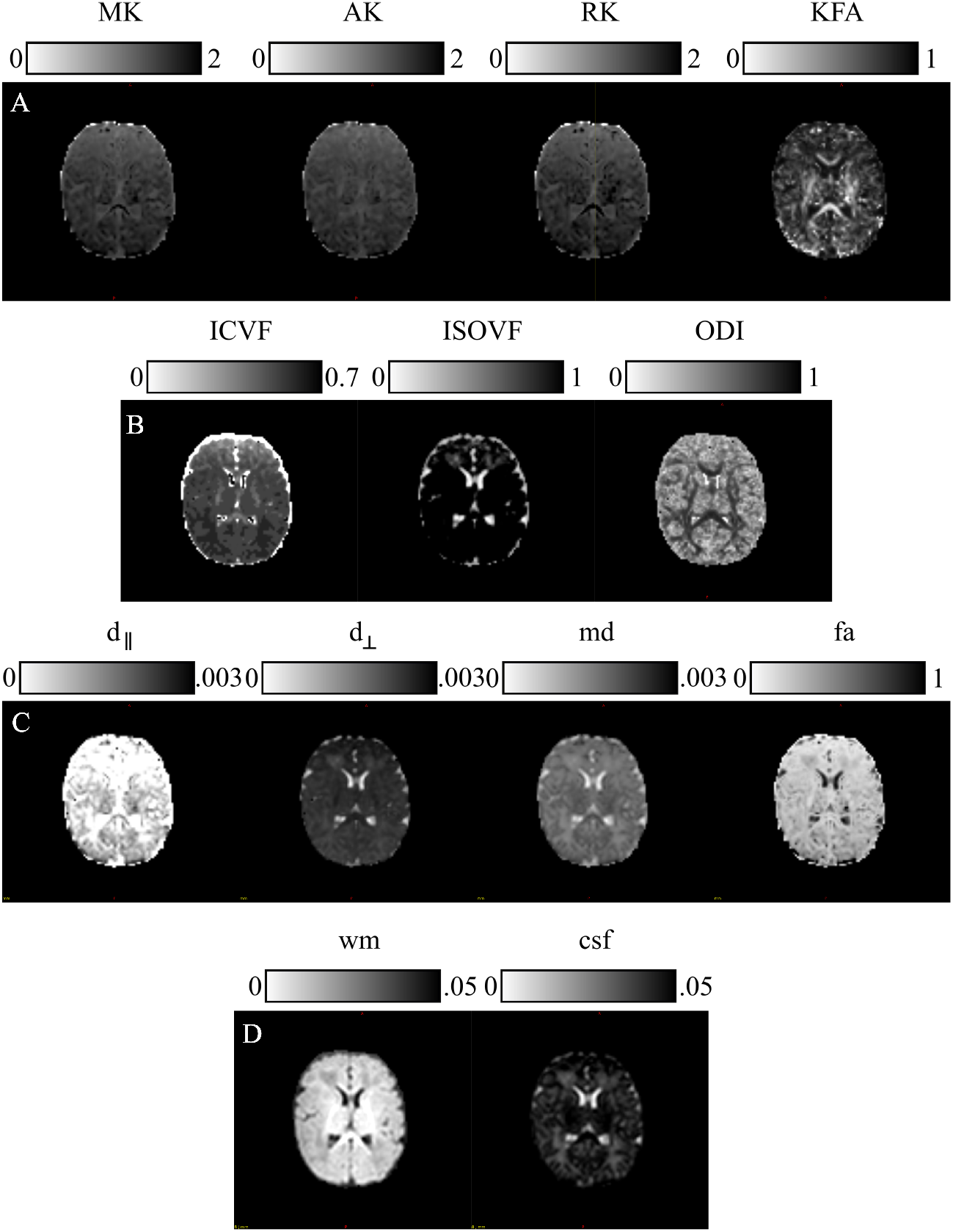
Microstructural Models: parametric scalar maps derived from all the HARDI models employed for this study: **(A)** Diffusion Kurtosis Imaging (DKI), **(B)** Neurite Orientation Dispersion and Density imaging (NODDI), **(C)** Fiber Orientation Estimated using Continuous Axially Symmetric Tensors (FORECAST), **(D)** Multi-Shell Multi-Tissue Constrained Spherical Deconvolution (MSMT CSD).

###### Diffusion Kurtosis Imaging

We performed the estimation of diffusion maps through DIPY v.1.4.0, (https://dipy.org) (Garyfallidis et al., 2014). Standard DKI parametric maps - Mean Kurtosis (MK), Axial Kurtosis (AK), Radial Kurtosis (RK), and Kurtosis Fractional Anisotropy (KFA) - were thus generated. Since these measures are susceptible to high amplitude outliers, we removed their impact by limiting metrics’ extraction within the typical range (0,3).

###### Neurite Orientation Dispersion and Density imaging

We computed NODDI-related measures - Intra Cellular Volume Fraction (ICVF), ISOtropic Volume Fraction (ISOVF), and Orientation Dispersion Index (ODI) - with a linear framework for Accelerated Microstructure Imaging via Convex Optimization (AMICO) implemented in Python (https://github.com/daducci/AMICO), which, through a convex optimization approach, drastically accelerates the fit of advanced dMRI techniques while preserving accuracy and precision in the estimated parameters, thus meeting real application demands (Daducci et al., 2015).

###### Fiber Orientation Estimated using Continuous Axially Symmetric Tensors

We resorted to DIPY also for the computation of measures derived from the FORECAST model). We used 6 as spherical harmonics order (*sh_order*) for the fibre Orientation Distribution Function (fODF) and CSD as spherical deconvolution algorithm for the FORECAST basis fitting (*dec_alg*) to extract crossing invariant tensor indices.

###### Multi-Shell Multi-Tissue Constrained Spherical Deconvolution

Application of MSMT CSD has been performed in MRtrix3 (http://www.mrtrix.org/). For response function estimation, used as the kernel by the deconvolution algorithm, we resorted to *dhollander* approach, suitable for computing MSMT response functions in case of multi-tissue variants of SD and more reliable in case of neonates (Dhollander et al., 2016, 2019). However, given the poor WM/GM contrast inherent to neonatal scans (Dhollander et al., 2018), we were limited to extracting tissue-specific ODF just for WM and CSF. Moreover, since interested in performing population studies, we used the same response function for all our cohorts. To this end, we calculated the average tissue response function for all our subjects exclusively for WM and CSF responses.

#### Post processing

##### Tract-Based Spatial Statistics

###### FA skeleton generation

We first used TBSS, the most popular voxel-wise statistical inference for WM anatomy (Bach et al., 2014), to inspect potential per-voxel differences across microstructural derived markers typical of preterm birth compared to term-born controls. However, once again neonatal imaging caused the standard TBSS pipeline developed in FSL (https://fsl.fmrib.ox.ac.uk/fsl/fslwiki to present technical challenges, due to smaller anatomical dimension and lower image contrast and resolution. We thus integrated it with DTI-TK (http://dti-tk.sourceforge.net/pmwiki/pmwiki.php?n=Documentation.TBSS)), as suggested also in (Bach et al., 2014; Tokariev et al., 2020).

The latter is a spatial normalization and atlas construction toolkit optimized for examining WM morphometry resorting to tensor-based registration able to leverage rich discriminating features afforded by DTI.

As a first step, we computed DTI tensor for each subject - through FSL, given DTI-TK interoperability with FSL - limiting to the *b*=700 s/mm^2^ shell rather than the whole multi-shell diffusion volume. Indeed, this was the alternative producing the best outcome in later analysis steps (e.g., TBSS FA skeleton generation). After quality control of DT images, we then moved to the registration and spatial normalization of DTI volumes, which is the core functionality of DTI-TK.

We opted for bootstrapping a population-specific DTI template from our cohort of study without relying on an existing template, given the age range under analysis. To this end, we employed all datasets to build our ad-hoc template in order to better capture within-population features (Supplementary Figure S1). Hence, we performed tensor-based registration of each subject to the template space, through consecutive steps aimed at iteratively refining the template and improving the registration outcome.

Then, we (i) generated the FA map of the high-resolution population-specific DTI template; (ii) resorted to FSL default command `tbss_skeleton` for the resulting WM skeleton; (iii) produced each FA map of the spatially normalized high-resolution DTI data, merging them together (`fslmerge` in FSL) to form a 4D FA map and its corresponding binary mask.

From this point on we thus moved to the default procedure in FSL with the last two steps, that is `tbss_4_prestats` and stats (`randomise`). The first one aims at thresholding the mean FA skeleton image at a suitable level. The latter is chosen by visual inspection, setting a value able to retain common major WM tracts avoiding those subject to excessive cross-subject variability and where the nonlinear registration has not been able to attain good alignments - 0.1 in our case, consistent with other works on neonates (Ball et al., 2010). Later, it projects all subjects’ FA data onto the resulting mean FA skeleton. This 4D image file containing the projected skeletonised FA data is thus fed into voxelwise statistics. The second one is aimed at finding voxels which significantly differ between our two cohorts in comparison. To this end, it applies cluster-size thresholding to the data, in which the size of the cluster is determined by 500 permutations by using Randomise FSL’s tool (http://fsl.fmrib.ox.ac.uk/fsl/fslwiki/Randomise). A threshold of *p* < 0.05 (95^th^ percentile of the distribution) is set for the clusters, corrected for multiple comparisons across space.

###### Non FA metrics

In order to extend TBSS analysis to diffusion-derived measures other than FA, we repeated DTI-TK + TBSS steps with some variations, similar to what was done in (Timmers et al., 2016). We thus converted each microstructural scalar map to the DTI-TK coordinates. Then, we reapplied to each measure the nonlinear registration transform previously obtained to transfer each FA map to the population-specific template. This procedure was repeated for each of the microstructural measures analysed in the current work.

##### Predictive model

###### Machine Learning methods for classification

Moving to ML analysis, we performed preterm/term-born subject classification based on a predictive model. To prevent overfitting, given the small amount of data available to train our model, we thus resorted to an SVM framework to categorize preterm-born and term-born individuals based on whole-brain WM skeleton estimated using TBSS. Indeed, among the variety of ML techniques applied so far in neuroimaging settings, SVM has emerged as one of the most popular ML methods (Chin et al., 2018; Chu et al., 2015), able to effectively cope with high-dimensional data and provide good classification results (Vapnik, 1999). We also carried out a further analysis to investigate how the performance changes by varying the input dimension of our data through feature selection and, then, we trained a classification model based on related findings. For the implementation of ML methods, we resorted to scikit-learn free software in Python (https://scikit-learn.org/stable/).

###### Experimental design

The experiments we carried out can be subdivided into two phases (Figure 4).

**Figure 3.**
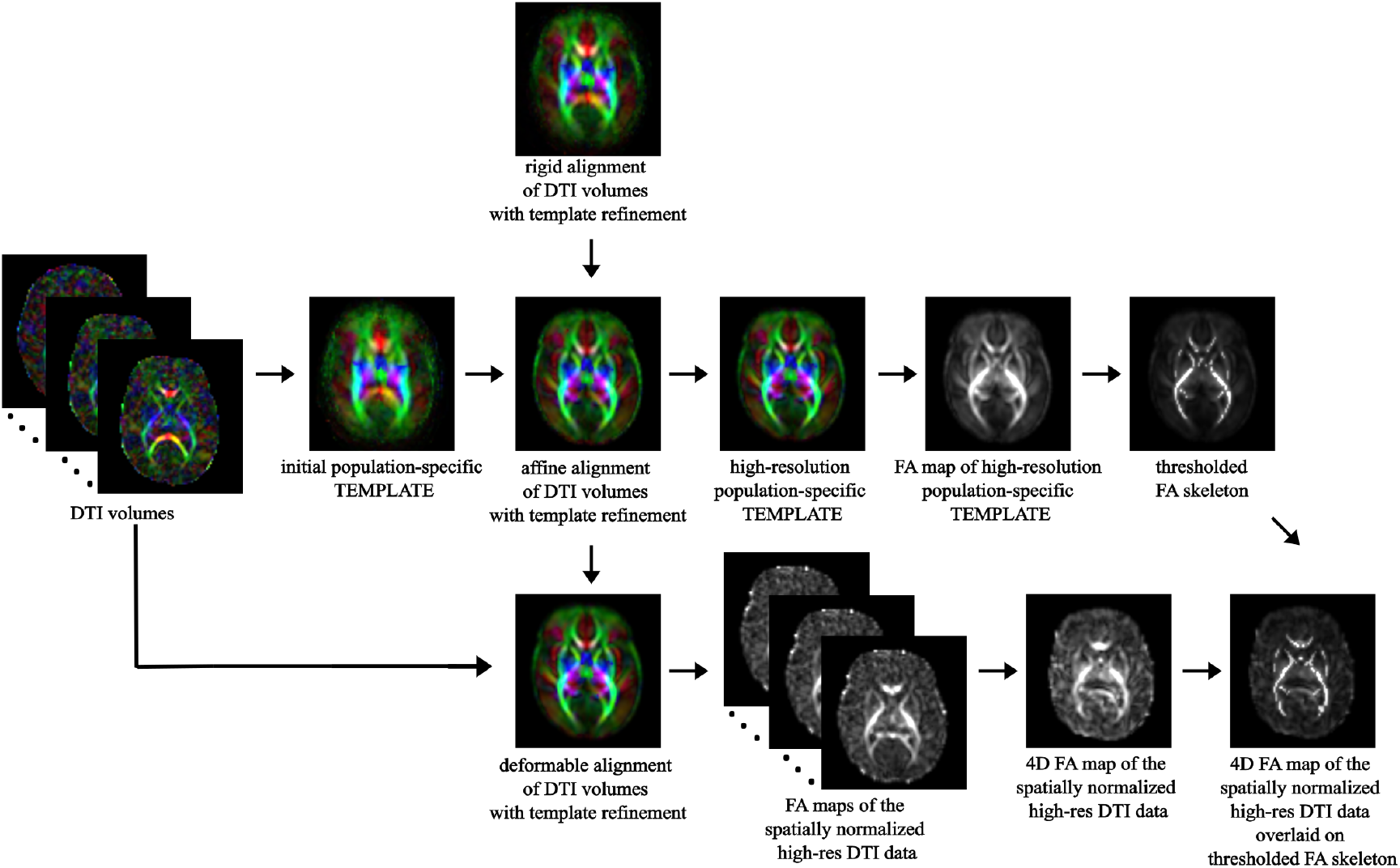
TBSS pipeline: overview of the main steps of TBSS framework, from spatial normalization of DTI volumes to bootstrapping the within-population template, to skeletonisation of the template’s FA map and projection of each subject’s FA onto the skeleton.

**Figure 4.**
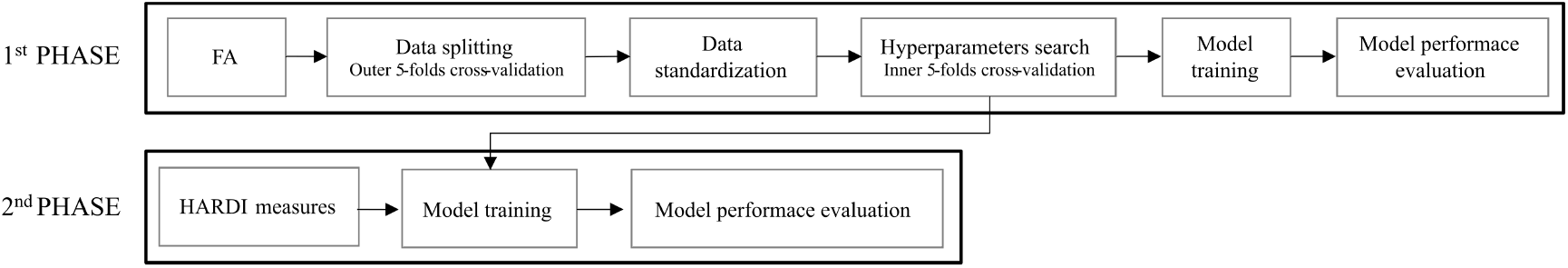
Experimental design for SVM classification: in a first phase a SVM classification estimator is chosen to best perform on FA skeletonised data; in a second phase the selected model is extended to other non-FA measures.

In the first phase, we adopted SVM to perform binary classification starting with the FA map warped to common TBSS space and masked by thresholded WM skeleton for all 69 infants involved. We then split the dataset into learning and testing by stratified 5-fold cross-validation (*outer-CV*) in order to increase the numerosity of our dataset while preserving the same class ratio throughout the *K* folds as the ratio in the original dataset. For each fold, we thus applied data standardization both on the learning set and the test set. We then further split the learning set into training and validation sets, named *inner-CV*, for exhaustively tuning the model hyperparameters with the GridSearchCV instance. We thus looked for the best hyperparameter grid by choosing the one that produced the lowest prediction error. This set included: (i) the best penalty term C (among 0.001,0.01,0.1,1,10,100,10^6^); (ii) the best kernel (among linear, radial basis function and polynomial with default degree=3) and (iii) the optimal number of features (selecting 20%, 40%, 60%, 80% and 100% of the input dataset with SelectKBest method). For each combination of hyperparameters, we fitted a model on the training set and thus evaluated its performance by computing the average f1-score across folds on the validation set. By selecting the set of parameters whose average f1-score was the best, we then trained such an SVM model on the learning set and subsequently evaluated its performance in terms of average and standard deviation of accuracy, precision, recall, f1-score, and Receiver Operating Characteristic (ROC) across folds on the unseen test set.

In the second phase, once selected the model classifier offering the best performance on FA data, we further evaluated the classification performance when giving as inputs the parametric measures from other microstructural models than DTI. In this phase, we did not perform any *inner-CV* as we did not introduce a hyperparameters search. Conversely, for each input variable, we again carried out the *outer*-*CV* to provide a more robust evaluation of the model. We thus trained the model on the learning set and then assessed the model on the test set computing the average and standard deviation of usual scores.

##### Weight maps extraction and comparison with TBSS

Finally, in order to relate the results from TBSS with those coming from ML, we extracted weight maps from the selected SVM classifier within *outer-CV*, averaged them across the 5 folds, normalized between 0 and 1 and reshaped them as the input 3D TBSS skeleton for mere visual comparison. The weights are SVM coefficients determining the discriminant hyperplane, which depicts the relevance of each voxel for the classification between positive and negative conditions.

We thus computed standard Pearson’s correlation between normalized SVM weight maps and TBSS normalized significance maps (p-maps) for each of the microstructural measures analyzed. In order to further inspect the overlap between WM discriminating regions detected by ML and TBSS, we related Pearson’s correlation with Wasserstein Distance (WD) metric, so as to quantify the distance between the two distributions.

##### Canonical Correlation Analysis

The CCA method is based on establishing linear relationships between two or more sets of variables, to find out the inter-subject co-variances. CCA looks for two or more sets of transformed variates - Canonical Components (CC) or Variates (CV) - to assume maximum correlation across the two datasets, while being uncorrelated within each dataset. Details about its mathematical formulation are illustrated in Appendix A

In our study, we resorted to the open-source Python package Pyrcca (Bilenko and Gallant, 2016) to perform a multi-set CCA based on fusing together all advanced dMRI models under analysis (Figure S2). We used as input all 14 HARDI features, after filling in missing values and z-scoring. To reduce the computational complexity of the analysis, a linear kernel was used. Moreover, we opted for a regularized kernel CCA to avoid overfitting, given the low numerosity of our datasets and to relax the orthogonality constraint between the canonical components. Finally, we estimated the optimal set of CCA hyperparameters - the regularization coefficient and the number of canonical components - empirically by using grid search with cross-validation. Specifically, the optimal regularization parameter was chosen from a logarithmically spaced range of 10 values between 1 × 10 ^−4^ and 1 × 10^2^; while the optimal number of components was chosen between 1 and 5. We selected these ranges based on pilot analyses performed on an independent dataset that was not used for this publication.

###### Shared/Distinct Abnormalities

As in (Sui et al., 2013), we inspected group differences between the two cohorts by performing a non-parametric Mann-Whitney U Test between each pair of Canonical Components, to look for the variants showing abnormalities associated with preterm birth. The statistical survey was followed by the Benjamini-Hochberg correction method for multiple comparisons (Benjamini and Hochberg, 1995). If the components show group differences in more than one modality, they are called modality-common group-discriminative CVs. Conversely, if the components show group differences only in a single modality, they are called modality-unique group-discriminative CVs.

###### Inter-modality correlation

We then investigated the inter-correlation existing between microstructural dMRI models, by looking at the Canonical Correlation Coefficients (CCC), with a view to establishing whether the joint-group discriminative components additionally have strong inter-modality correlation, which would reflect the interaction and correspondence among modalities.

## RESULTS

### Voxel-wise statistics on the WM skeletonised data

Cross-subject voxel-wise statistics unravelled significantly different voxels exclusively on a subset of the microstructural maps under consideration.

Specifically, compared with the term cohort, the preterm group showed a significant decrease in Fractional Anisotropy (FA), Mean Kurtosis (MK), Axial Kurtosis (AK), IntraCellular Volume Fraction (ICVF) and fractional anisotropy (fa). The WM regions with significant between-group differences in diffusion metrics are shown in Figure 5. Conversely, no significant differences have been captured by TBSS analysis in Radial Kurtosis (RK), Kurtosis Fractional Anisotropy (KFA), ISOtropic Volume Fraction (ISOVF), Orientation Dispersion (OD), mean diffusivity (MD), parallel diffusivity (d_∥_), perpendicular diffusivity (d_⊥_) either in MSMT-derived measures.

**Figure 5.**
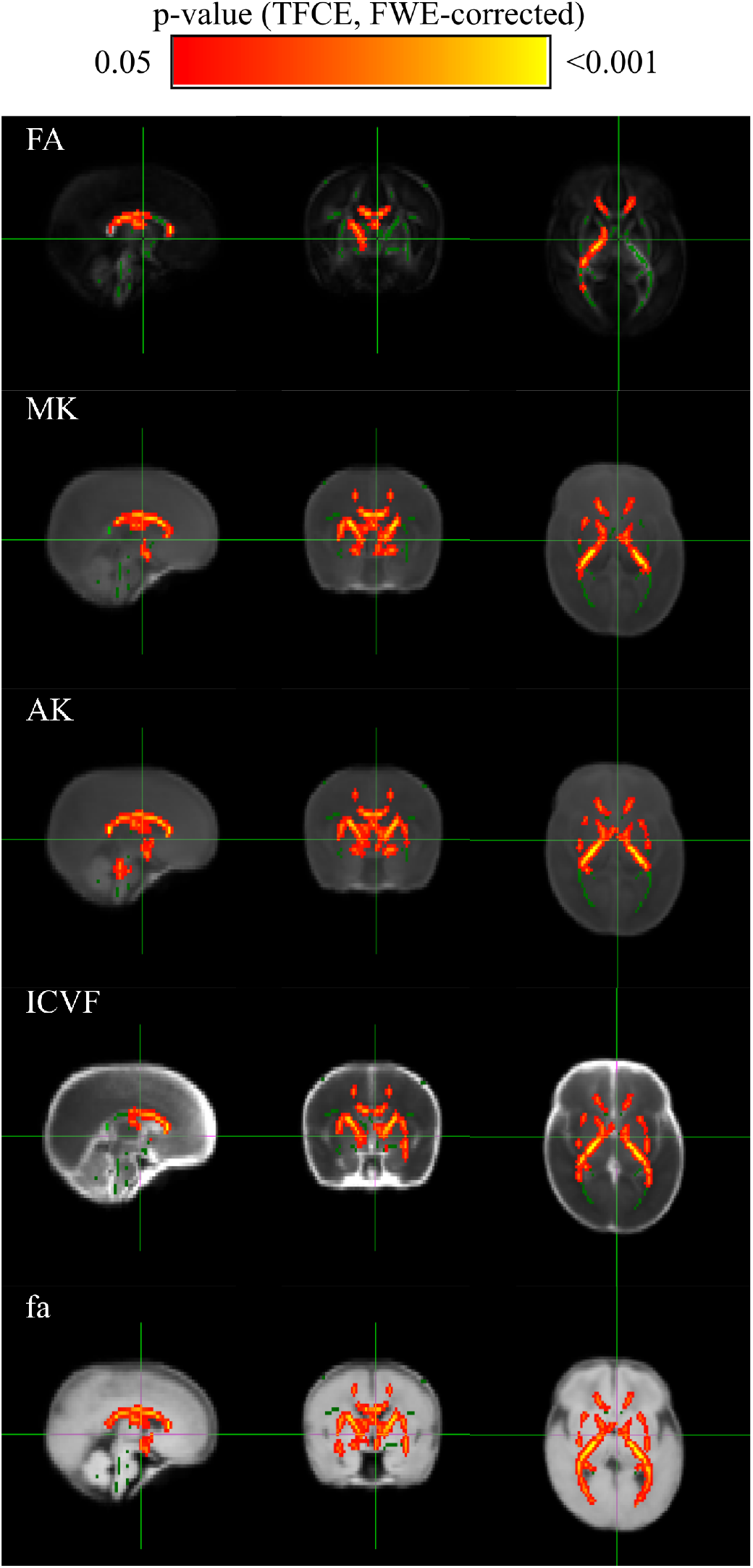
TBSS results: Group-level voxel-wise statistical difference maps for Fractional Anisotropy (FA), Mean Kurtosis (MK), Axial Kurtosis (AK), IntraCellular Volume Fraction (ICVF) and fractional anisotropy (fa) between preterm and term-born cohort. Green indicates the FA skeleton with a threshold of 0.1, which highlights the tracts used in the comparison. Red-Yellow indicates the regions with decreased metrics’ values in the preterm group, after a non-parametric 2-sample t-test with Family-Wise Error (FWE)-corrected *p*-values using Threshold-Free Cluster Enhancement (TCFE).

More in detail, compared with the term group, the preterm cohort had significantly decreased FA values in widespread WM areas predominately in the genu, body and splenium of the corpus callosum, right internal capsule, corona radiata, and posterior thalamic radiation. The distribution of areas with decreased MK was similar but extending bilaterally with respect to the areas with decreased FA and including also right external capsule. AK exhibited a pattern analogous to MK whilst comprising bilateral external capsule. The same applies for ICVF metric except for excluding the genu of the corpus callosum. The amount of WM areas showing a significant decrease in prematurity increased for fa parameter derived from FORECAST model, which extended to the whole corpus callosum, bilateral internal capsule, external capsule and anterior corona radiata and, finally, posterior thalamic radiation (including the optic radiation).

### SVM classification on the skeletonised data

Since the performance of a model significantly depends on the value of its hyperparameters, first of all, we focused on hyperparameter tuning in order to determine the optimal values for our classification estimator.

In this respect, Figure 6A shows the result of cross-validated grid-search over the parameter grid across each of the 5 folds. Furthermore, based on the selected hyperparameters, we fitted our model on the training set and evaluated its performance on the test set in terms of f1-score, accuracy, precision, recall and ROC across each of the 5-fold (Figure 6B). In order to establish the best estimator possible based on the input data, we counted into how many folds a variable was selected and could thus conclude that penalty term *C* and *linear* kernel were the most frequently selected hyperparameters. Conversely, the search turned out to be less stable in terms of the optimal number of features, which varied at every fold (Figure 6C). Therefore, in order to set the last missing parameter for our estimator, we set *C* and kernel according their most chosen values while varying the number of features as a percentage of the total amount.

**Figure 6.**
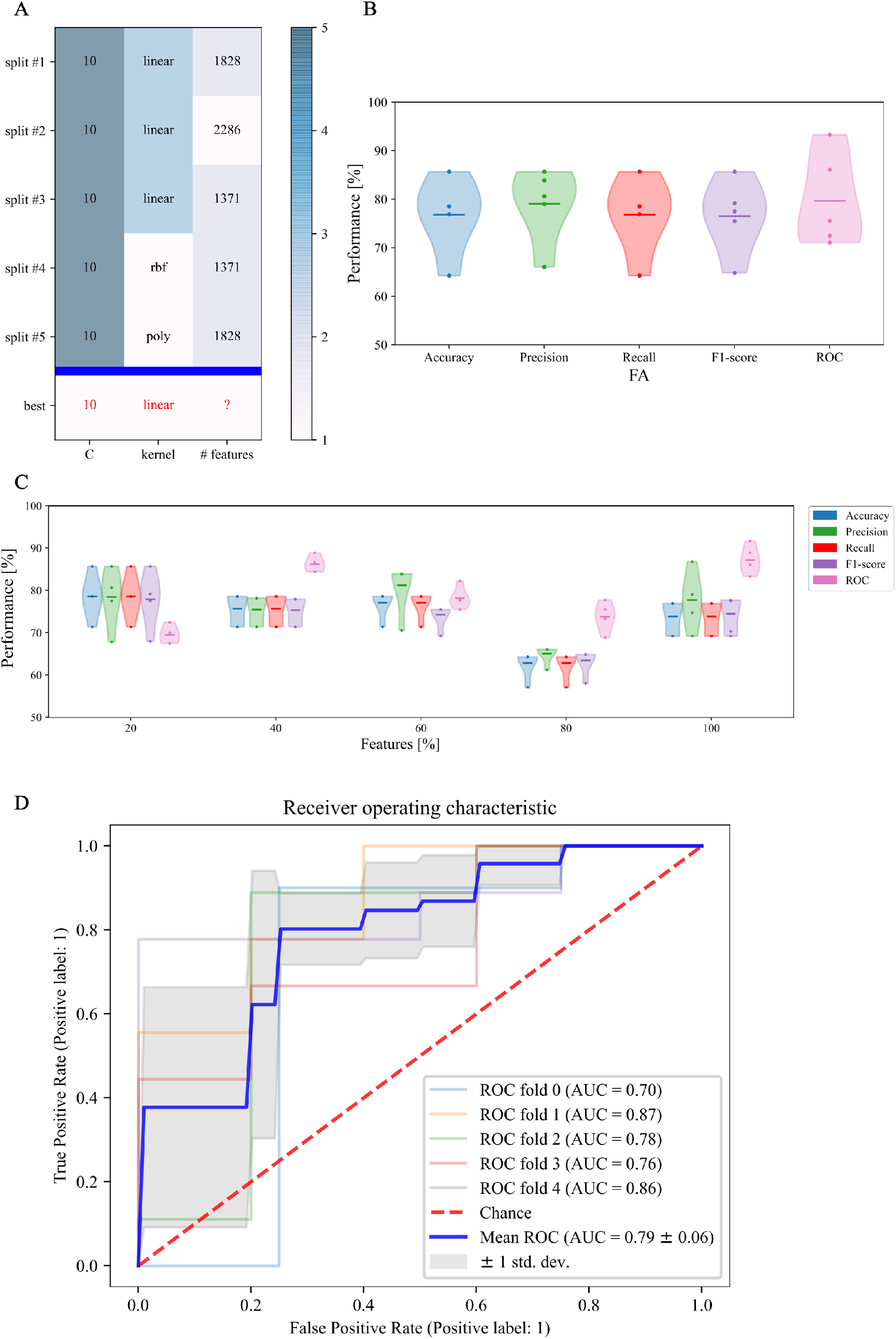
First phase: SVM training on FA skeletonised data: **(A)** cross-validated search of the best set of hyperparameters for our SVM estimator on stratified 5-fold data; **(B)** relative performance for every score across folds; **(C)** Different sets of selected features along with relative performance for every score across folds; and **(D)** area under the curve score

Figure 6C confirms that in our case feature selection is not beneficial for improving the classification performance. Indeed, both average value and standard deviation across folds of each score remain constant with variable subset of features. In addition, average ROC score proves to be maximal (0.87%) when including the whole feature amount. We thus opted for avoiding feature reduction and kept the whole of features to define the final version of our SVM estimator. As regards this definitive version of the classifier, a detailed plot of the ROC curve profile for every fold is displayed in Figure 6D. We subsequently trained a classification model without hyperparameters’ search (*inner-CV*) using as input variables the features derived from other microstructural HARDI models. Performance in terms of f1-score, accuracy, precision, recall and ROC for the whole set of microstructural parameters, including FA, is reported in Figure 7. Of note, among the whole set of features, the ones exhibiting highest discriminative power in terms of SVM classification are those probing in general fibers’ anisotropy and directionality, namely FA, KFA, OD and fa, for which all scores overcome 75%, 74%, 70% and 74% levels on average, respectively.

**Figure 7.**
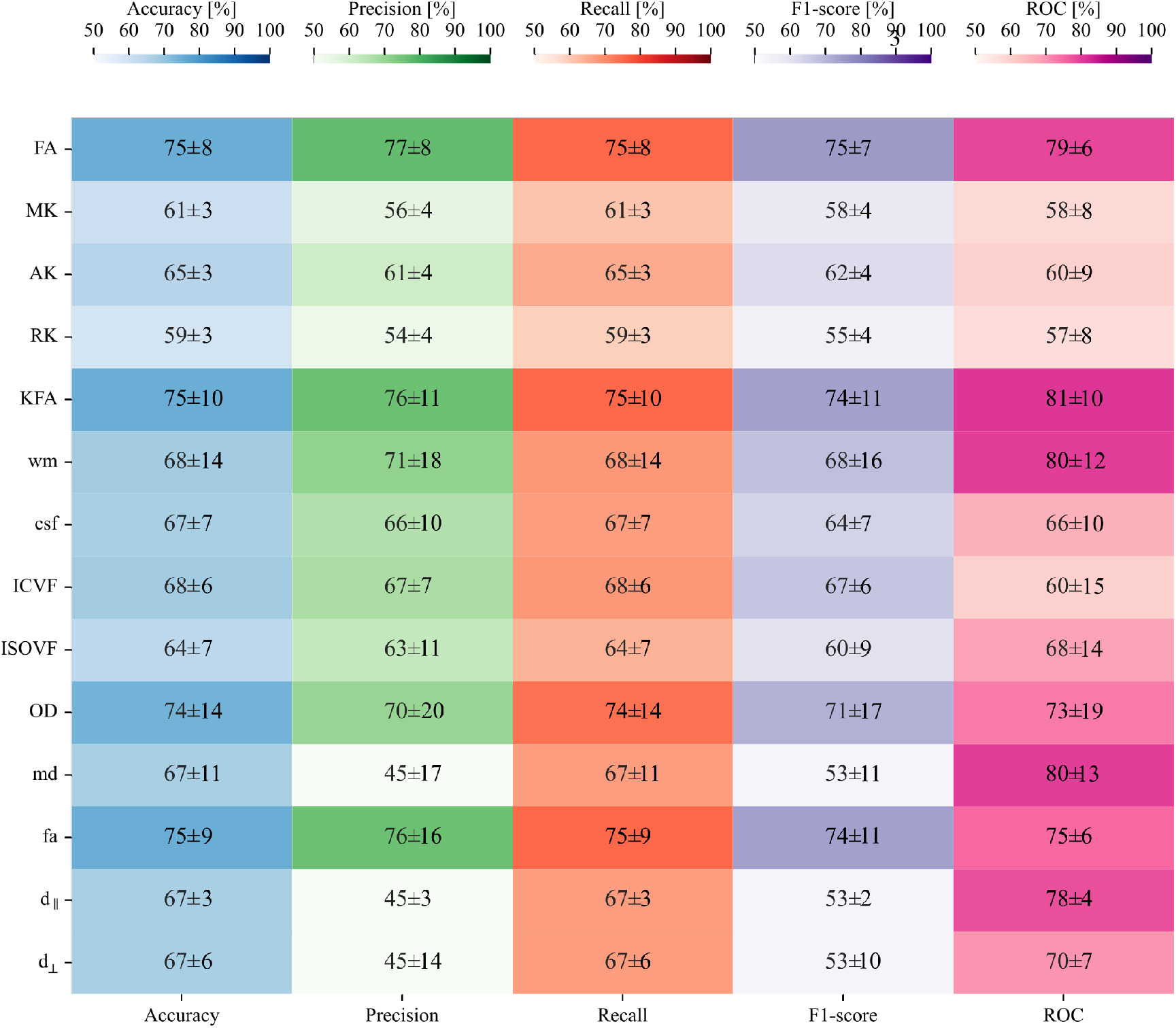
Second phase: SVM testing on non-FA skeletonised data: heatmap containing average and relative standard deviation, in percentage, of each score and for all the HARDI measures under analysis.

### Comparison between SVM and TBSS approach

Relating TBSS significance map with weights extracted from linear SVM showed statistically significant Pearson’s correlation for all microstructural measures considered (*p* < 10 ^-2^) (see Table 3). This relationship was further confirmed by inspecting the association between Pearson’s correlation coefficient and WD, reported in Figure 8, showing a trend of direct proportionality. Indeed, those measures exhibiting a higher absolute correlation also presented a lower WD, thus confirming similarity between the two methods. Correlation was high (*r* = 0.61) for d_∥_ parameter, intermediate (*r* ∈ 0.45 – 0.51) for RK, KFA, FA and OD, moderate (*r* ∈ 0.28 – 0.35) for MK, AK, MD, fa and ICVF and low (*r* ∈ 0.05 – 0.14) for d_⊥_, CSD-related measures and ISOVF. These results suggest an overall good, though the measure-dependent rate of agreement between p-maps derived by TBSS approach and weights probing discriminative power of SVM.

**Table 3.** Comparison between TBSS voxelwise statistics and SVM classification

| | Pearson's $r$ | $p$-value | Wasserstein distance |
| --- | --- | --- | --- |
| **FA** | -0.45 | <0.0001 | 0.32 |
| **MK** | -0.34 | <0.0001 | 0.43 |
| **AK** | -0.33 | <0.0001 | 0.41 |
| **RK** | -0.46 | <0.0001 | 0.36 |
| **KFA** | -0.48 | <0.0001 | 0.35 |
| **$d_{\parallel}$** | -0.61 | <0.0001 | 0.20 |
| **$d_{\perp}$** | -0.14 | <0.0001 | 0.48 |
| **md** | -0.33 | <0.0001 | 0.36 |
| **fa** | -0.28 | <0.0001 | 0.40 |
| **wm** | -0.14 | <0.0001 | 0.42 |
| **csf** | -0.11 | <0.0001 | 0.55 |
| **ICVF** | -0.35 | <0.0001 | 0.36 |
| **ODI** | -0.51 | <0.0001 | 0.41 |
| **ISOVF** | -0.05 | 0.013 | 0.46 |

**Figure 8.**
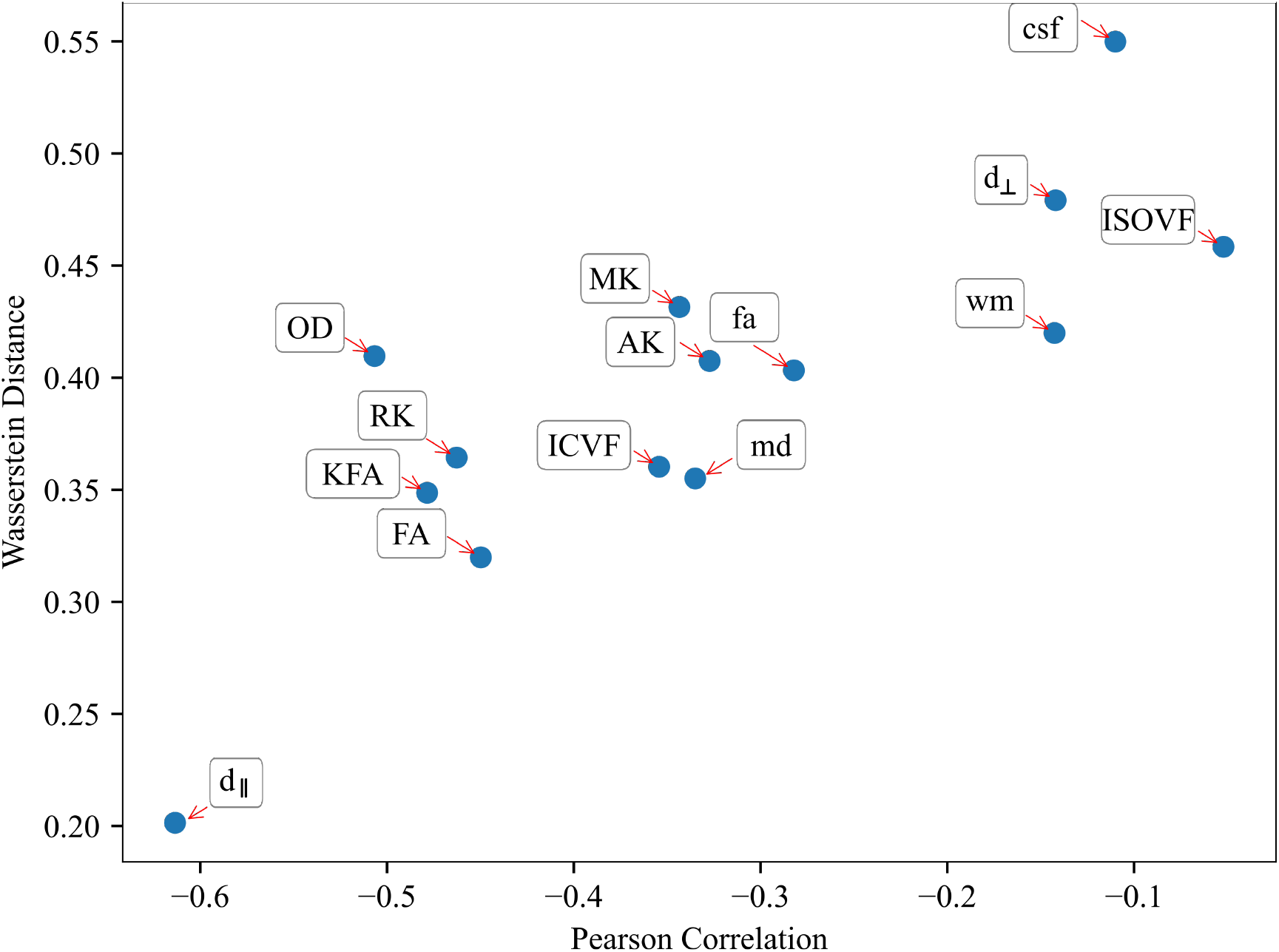
Relationship between Pearson’s correlation and Wasserstein Distance for HARDI microstructural models: the two measures show a good trend of association throughout all HARDI parameters considered.

### Canonical Correlation Analysis

Moving to CCA analysis, Pyrcca cross-validated hyperparameters search detected the optimal regularization coefficient equal to 0.01, and the optimal number of CVs to 4. Preliminarily, the results of CCA analysis were evaluated in terms of Canonical Correlations to determine the number of meaningful Canonical Components recovered by Pyrcca. Figure 9 contains a heatmap of pair-wise correlations between the 14 HARDI measures for each of the 4 sets of Canonical Components. CCA analysis applied to our cohort, from Mann-Whitney U Test, unravelled group differences in the 4^th^ Canonical Component, for which statistically significant differences between preterm and term subjects have been found in d_∥_ and ISOVF even after FDR correction (*p* < 0.05), thus being a modality common or joint group discriminative independent component. This is depicted in Figure 10A, with violin plots of CVs having statistically significant differences between preterm and term-born subjects. Interestingly, the intramodal connection within the joint-discriminative independent component (4^th^) indicates a high correlation (*r* = 0.62). (see Figure 9). Furthermore, to visually mark out detected differences between the two groups, we displayed derived spatial maps only for the specific joint 328 group-differentiative canonical component. In Figure 10B each input feature transformed into a z-score is reported to highlight statistically significant group-discriminating subsets of voxels.

**Figure 9.**
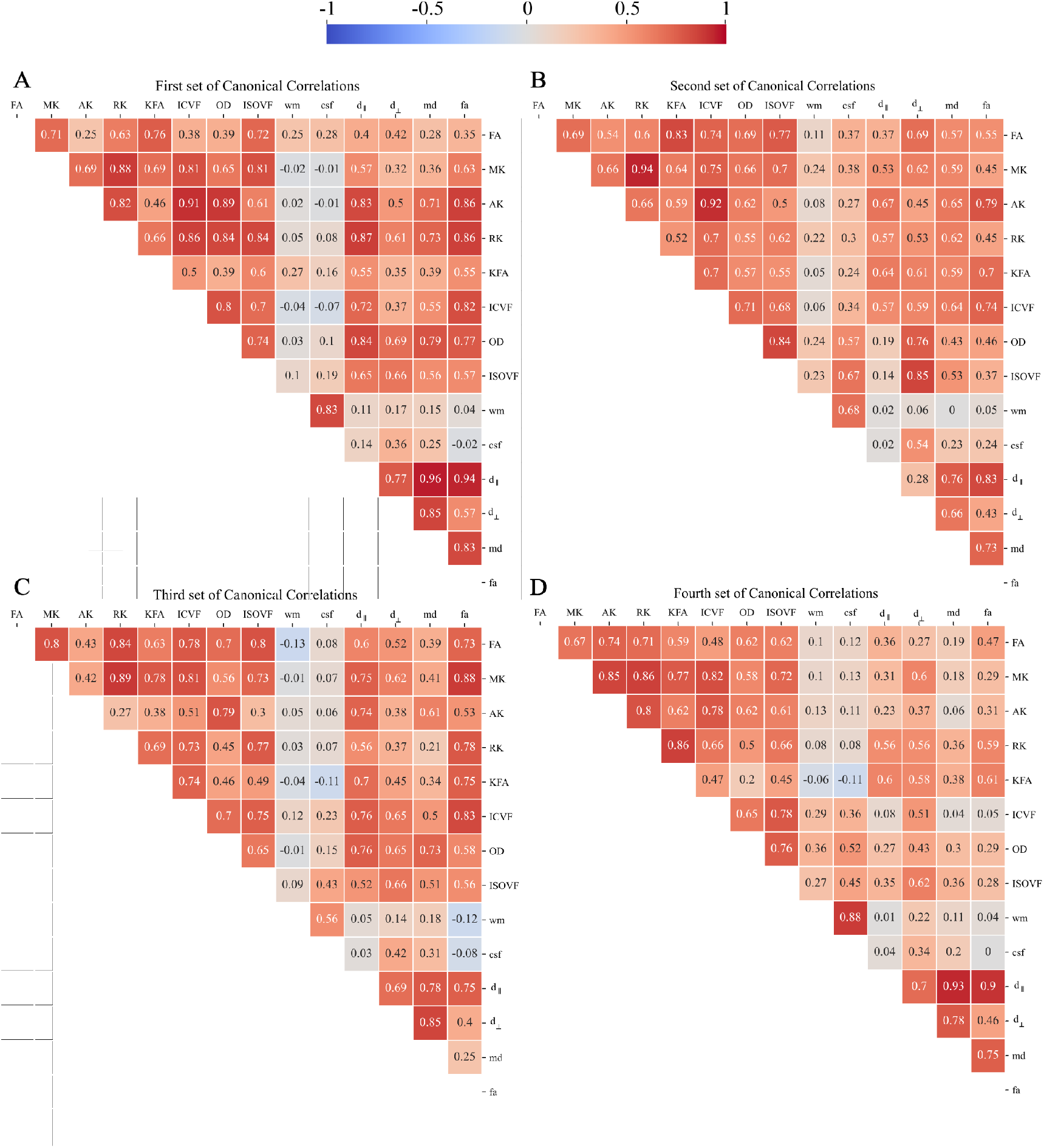
Pair-wise Canonical Correlation Matrices from CCA analysis for: **(A)** first Canonical Component, **(B)** second Canonical Component, **(C)** third Canonical Component, and **(D)** fourth Canonical Component.

**Figure 10.**
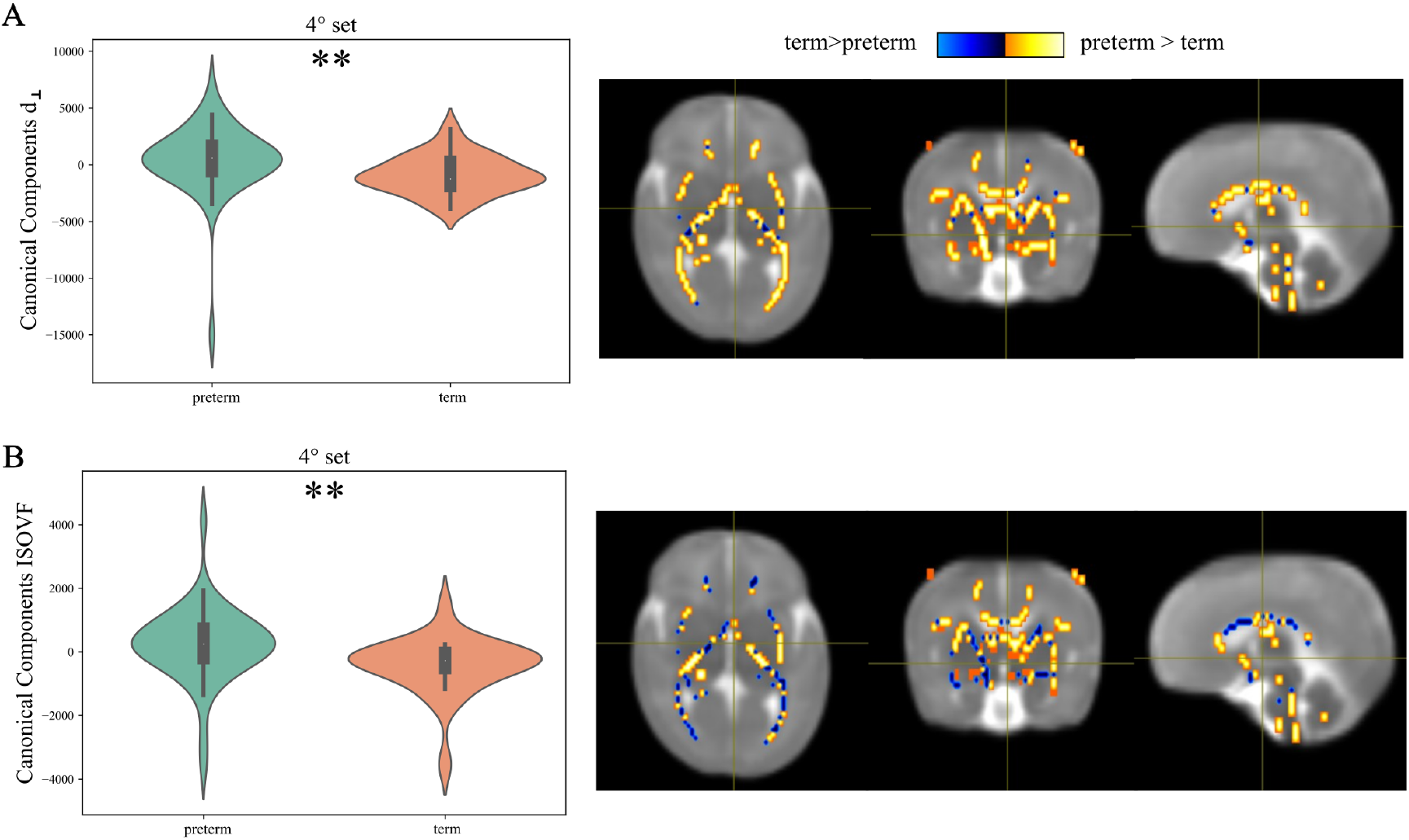
Joint group-discriminative component for d_⊥_ and ISOVF modalities: **(A)** Violin plots of the loading parameters for 4^th^ component for each modality, with the *p* values of Mann Whitney U Test between term and preterm participants; **(B)** Group-discriminating regions across all modalities. The Z-scored spatial maps exhibit positive Z-values (orange regions) meaning preterm > term subjects, and the negative Z-values (blue regions) mean term > preterm.

## DISCUSSION

In this work, based on observations that integrating data from different MRI sequences enhances anatomic characterization (Ball et al., 2017; Kulikova et al., 2015; Thompson et al., 2019a), we aimed at combining the most popular HARDI-based microstructural models to supplement the understanding of WM differentiation in the preterm cohort for an early clinical diagnosis. Indeed, HARDI acquisitions potentially allow to investigate of neurodevelopment with a higher degree of sensitivity and earliness, yet at the price of pitfalls in scanning vulnerable preterm infants. Of note, obtaining good quality images is particularly challenging in this patient group primarily as motion-sensitive and artefact-prone (Jones and Cercignani, 2010).

The first achievement of the present work is thus the successful implementation of the overall image processing pipeline able to fully investigate our cohort. This was possible thanks to the setting of a suitable acquisition protocol, fine-tuned to the specific characteristic of infant’s brain, and supported by the patient’s mild sedation, monitoring and fixation and resulting in high-quality images suitable for HARDI modelling. Another crucial role is played by preprocessing: a dedicated workflow, whose denoising is confirmed as an essential step, is here key for subsequent analysis.

Moving to actual post-processing, the first tool we considered to investigate the fingerprint of preterm subjects was TBSS. In fact, the analytic method of TBSS offers a number of advantages over hypothesis-directed ROI analyses, in that it describes changes in WM microstructure in a 3D image space. Furthermore, resorting to a population-specific template ensured good alignment, essential for subsequent analyses, given the vast variability of brain morphology at these early stages of development. Through TBSS, we demonstrated that both FA and non-FA values can be useful measures to distinguish relevant WM tracts in preterm-born neonates at TEA from term-born controls. It was particularly notable that there was a correspondence between the distribution of areas with decreased FA and non-FA measures, with an expansion of WM discriminating areas over the main tracts, especially in the case of beyond-DTI measures.

This agrees with existing findings in the literature claiming that: (i) WM maturation is associated with increasing axonal organisation, pre-myelination and myelination, which progressively restricts water diffusion perpendicular to the direction of the axonal fiber; (ii) since premature birth may lead to relatively slow brain development in premature infants, there are some brain regions that are less developed than the full-term infants. This includes CC, ALIC, PLIC, and, more generally, all tracts subject to early myelination whose metabolism is thus vigorous and the oxygen demand is high, which makes these metabolically active areas the first to be damaged in case of risk factors for preterm birth (Ling et al., 2013). For the DTI measures, lower FA has been found across the WM in preterm infants compared with term-born infants (Pecheva et al., 2018; Hüppi et al., 1998b; Anjari et al., 2007; Thompson et al., 2011), which correlated with increased prematurity (Ball et al., 2010; Partridge et al., 2004; Ouyang et al., 2019a). Furthermore, WM diffusion measures in preterm infants at TEA have been related to subsequent neurodevelopmental performance. Decreased FA - together with increased MD and RD - in the WM at TEA are associated with worsened motor, cognitive, and language performance in early childhood (Counsell et al., 2008; Barnett et al., 2018) as well as a visual function (Bassi et al., 2008; Groppo et al., 2014). In (Zhao et al., 2021), kurtosis-related parameters, especially MK, showed to sensitively reflect the brain maturity of premature infants. Decreased MK values were registered in the preterm cohort due to the decreased density of cells and axon membranes associated with impaired brain development.

Similarly, NODDI model has been applied to investigate WM and GM maturation in the preterm brain (Kimpton et al., 2021; Batalle et al., 2019, 2017;Eaton-Rosen et al., 2015), finding that ICVF increases in the WM with increasing maturation, mainly attributed to increasing axonal growth/density/packing/diameter or pre-myelination/myelination changes, rather than changes in axon coherence or geometry. Moreover, greater ICVF in childhood has been associated with better neurodevelopmental outcomes, IQ (Young et al., 2019; Kelly et al., 2016), visual motor integration (Young et al., 2019), motor/behavioural/emotional scores (Kelly et al., 2016), language (Mürner-Lavanchy et al., 2018) and maths (Collins et al., 2019).

Finally, although not previously investigated in the case of preterm subjects, FA parameter derived from the FORECAST model falls into those measures exhibiting significant differences from preterm to term-born infants, presumably for being the equivalent of the DTI FA yet far more sensitive to the underlying fiber microanatomy.

The second perspective from which we examined our cohort was an SVM-based approach aimed at a more individualized classification method to overcome shortcomings of group-wise investigations. The good achievement of the SVM in assigning a preterm-born or control individual to the correct group, based on a single MR image, indicates that the distinct brain development of preterm-born individuals can be successfully identified by ML methods. Indeed, considering the low sample size at disposal, much inferior to the number of features (i.e., image voxels), the SVM classifier managed to handle the issue of overfitting and proved a good performance both on FA skeletonised image, on which its model was designed, and on the vast majority of non-FA measures. Specifically, together with FA from DTI, other scalar parameters derived from DKI, NODDI and FORECAST exhibited good scores in terms both of f1/accuracy and, most importantly, of ROC - a significantly more meaningful measure of classifier performance than accuracy because it does not bias on size of test or evaluation data. Of note, preterm vs term classification accuracy achieved by the predictive model, however good, was not optimal. This may be due to the diffuse effect of preterm birth on WM microstructure, being optimally captured by methods not requiring anatomically constrained ROIs (Baykara et al., 2016;Blesa et al., 2020). Along with overall good performance scores, the selected classifier also showed strong robustness (i.e. limited variability across folds), another important indicator for model evaluation, assessing its stability. Furthermore, the evidence that most discriminating features in terms of SVM classification are related to fibers’ anisotropy stands for dismaturation or delay in myelination of WM tracts following preterm birth in contrast to term-born controls.

We then explored the relationship occurring between TBSS- and SVM-based methods, assessing the degree of overlap between the two survey methods. The observed negative Pearson’s correlation finds its explanation in considering that we compared a significance map from voxel-wise statistics made up of thresholded p-values and the map of SVM weight vectors serving as ranking metric for measuring feature importance (Gaonkar and Davatzikos, 2013). As a result, those voxels exhibiting a lower p-value correspondingly have a high ranking in the SVM model, which brings to the observed inverse trend. Agreement between voxel features for the two methods is just partial, to reflect the fact that a discrepancy actually exists between the two kinds of approach in voting WM connections with discriminative ability. Indeed, as already explained in the introduction of this work, ML approaches have entered neuroimaging field to try to improve data-driven extraction of knowledge about underlying biological correlates.

It is on the premise that integrating multiple datasets from the same participants can increase confidence when making conclusions to a greater degree than traditional statistical approaches (Sui et al., 2013) that we extended our investigation to considering simultaneously multiple microstructural models through CCA. In this paper, we investigated brain co-alterations from several advanced dMRI across preterm and term-born cohorts. To our knowledge, this is the first study to clarify preterm birth-related brain changes in different dMRI modalities via an intramodal data fusion model. Specifically, two further measures, d_⊥_ and ISOVF, emerged as relevant markers discriminative of preterm birth, other than those highlighted from TBSS or classification. This proves CCA’s capabilities to detect potentially hidden relationships between different imaging modalities beyond traditional methods, which could not be detected from a single dMRI model.

The brain regions exhibiting the strongest contributions to coherent changes related to preterm birth involve the majority of WM tracts detected with TBSS. More in detail, a simultaneous decrease in both d_⊥_ and ISOVF is observed in term-born group compared to the preterm one. Such findings are in line with previous studies (Vaher et al., 2022; Barnett et al., 2018; Pecheva et al., 2017; Thompson et al., 2019b) and consistent with the higher content of extracellular free-water expected in case of diffuse loss of WM microstructural integrity and organization inherent to preterm birth. Similarly, being d_⊥_ a more sensitive variant of DTI’s Radial Diffusivity, it proved in turn to be highly reflective of the lack of tortuosity imposed on water motion due to delayed development of the myelin sheath (Knight et al., 2018).

The overall results demonstrate that fusing information from different microstructural models even within the same imaging modality is able to provide a powerful and flexible tool to identify interesting associations between modalities as well as changes in these associations due to disease. Furthermore, these findings should be added to the body of literature suggesting generalised dysmaturation of the WM in preterm neonates.

We are aware that both voxel-wise statistical methods and, in particular, the ML approach benefits from large quantities of data. Consequently, one future step could be extending the current dataset in order to further improve our findings and, possibly, introducing a stratification based on the patient’s diagnoses. Indeed, we hypothesise that the greater potential of ML classification may be useful to distinguish specific patterns of WM tracts also between healthy and pathological subjects, not detectable with standard group-level methods. This is a possible interesting development of the current investigation, which could further highlight the contribution of ML-based approaches to neuroimaging.

## CONCLUSION

Results gathered so far from this study revealed that an intramodal dMRI approach can be a valuable tool to distinguish atypical brain microstructure at TEA when compared with a full-term group, regardless of the specific diagnosis based on radiological findings. This differentiation is attained at three different levels of investigations, in order to provide a more comprehensive, detailed and biologically meaningful interpretation of WM microstructure changes associated with prematurity. First, a classical group-level survey tool such as TBSS confirmed the increased sensitivity of advanced dMRI methods. Secondly, a state-of-the-art approach based on SVM classification achieved a good recognition rate. Furthermore, a comparison of the two methods showed a discrete agreement in selecting the most discriminating WM regions, mainly depending on the microstructural measure under consideration. Finally, CCA further represents a powerful tool to identify the inter-measure similarities between features associated with preterm birth in a data-driven way, without imposing an explicit model. Taken together, these findings suggest that exploiting synergy between modalities allows a more throughout investigation of the preterm birth phenomenon providing an unprecedented supplement to the understanding of biological mechanisms. Further work should focus on investigating how well these results generalize to data across centers and on what kind of improvements are needed to reach the end goal of predicting, on an individual basis, the specific outcome of subjects born preterm.

## ACKNOWLEDGMENTS

The authors would like to thank Prof. Luca Antonio Ramenghi [Neonatal Intensive Care Unit, IRCCS Istituto Giannina Gaslini, Genoa, Italy and Department of Neurosciences, Rehabilitation, Ophthalmology, Genetics, Maternal and Child Health (DINOGMI), University of Genoa, Italy] and the LIFT (Laboratorio di Imaging Funzionale 3 Tesla).

## A APPENDIX 1 Canonical Correlation Analysis

CCA aims at decomposing each input feature into a set of Canonical Variates *A_k_* and the corresponding Canonical Weigths *W_k_*, given by:

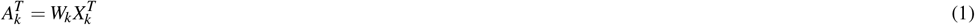

Where *X_k_* ∈ *R^N_x_V_k_^, A_k_* ∈ *R^NxD^, W_k_* ∈ *R^V_k_xD^*, *V_k_* is the number of variables in *X_k_, N* is the number of observations in *X_k_*, and D is the number of CV. From CCA, within an intramodal fusion approach, one can go backwards decomposing each input feature *X_k_* into a set of components Ck, and corresponding modulation profiles (inter-subject variations) *A_k_,* so that:

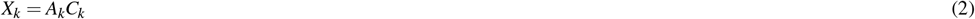

with:

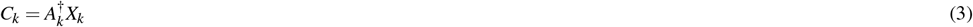

## B APPENDIX 2 Supplementary Material

**Figure S1.**
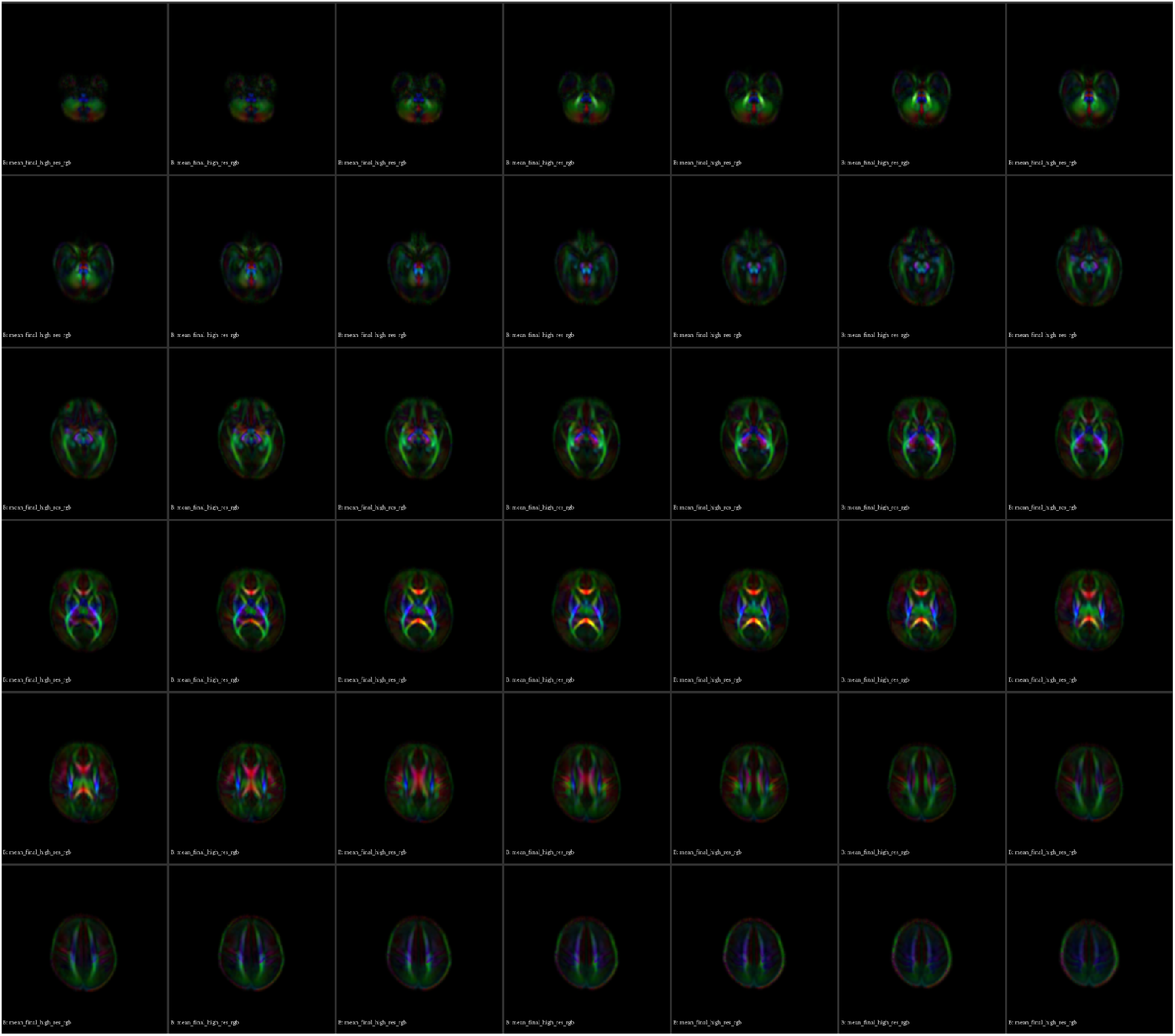
Population-specific DTI template: lightbox displaying axial views of the age-specific template created ad-hoc for performing normalization of DTI volumes within DTI-TK and subsequent WM skeleton creation within TBSS.

**Figure S2.**
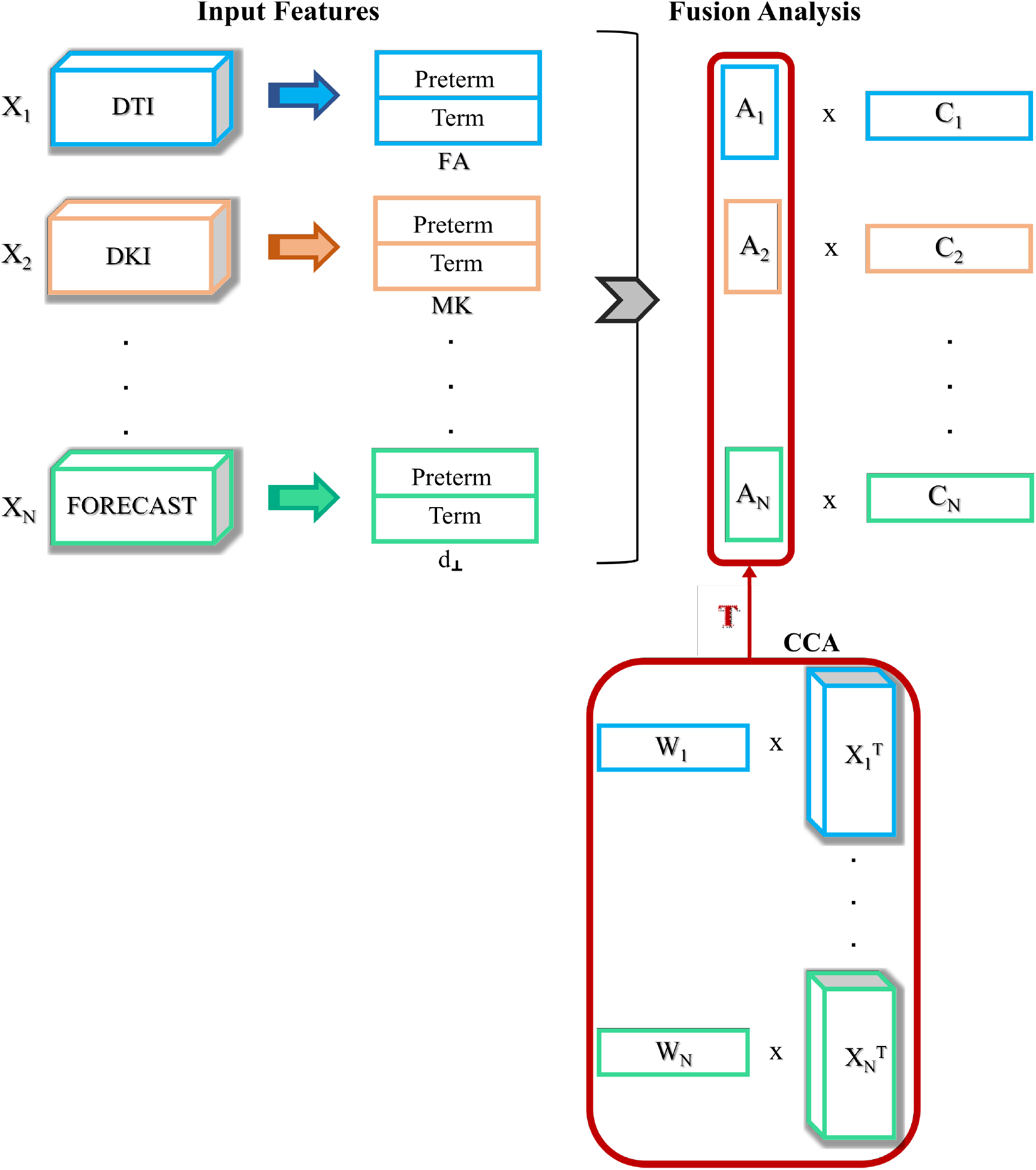
Canonical Correlation Analysis framework. applied to our intramodal dataset on the all the 14 HARDI microstructural measures.

## REFERENCES

Alexandrou, G., Mårtensson, G., Skiöld, B., Blennow, M., Ådén, U., and Vollmer, B. (2014). White matter microstructure is influenced by extremely preterm birth and neonatal respiratory factors. Acta paediatrica, 103(1):48–56.

Anderson, A. W. (2005). Measurement of fiber orientation distributions using high angular resolution diffusion imaging. Magnetic Resonance in Medicine: An Official Journal of the International Society for Magnetic Resonance in Medicine, 54(5):1194–1206.

Anjari, M., Srinivasan, L., Allsop, J. M., Hajnal, J. V., Rutherford, M. A., Edwards, A. D., and Counsell, S. J. (2007). Diffusion tensor imaging with tract-based spatial statistics reveals local white matter abnormalities in preterm infants. Neuroimage, 35(3):1021–1027.

Bach, M., Laun, F. B., Leemans, A., Tax, C. M., Biessels, G. J., Stieltjes, B., and Maier-Hein, K. H. (2014). Methodological considerations on tract-based spatial statistics (tbss). Neuroimage, 100:358–369.

Ball, G., Aljabar, P., Nongena, P., Kennea, N., Gonzalez-Cinca, N., Falconer, S., Chew, A. T., Harper, N., Wurie, J., Rutherford, M. A., et al. (2017). Multimodal image analysis of clinical influences on preterm brain development. Annals of neurology, 82(2):233–246.

Ball, G., Counsell, S. J., Anjari, M., Merchant, N., Arichi, T., Doria, V., Rutherford, M. A., Edwards, A. D., Rueckert, D., and Boardman, J. P. (2010). An optimised tract-based spatial statistics protocol for neonates: applications to prematurity and chronic lung disease. Neuroimage, 53(1):94–102.

Barnett, M. L., Tusor, N., Ball, G., Chew, A., Falconer, S., Aljabar, P., Kimpton, J. A., Kennea, N., Rutherford, M., Edwards, A. D., et al. (2018). Exploring the multiple-hit hypothesis of preterm white matter damage using diffusion mri. NeuroImage: Clinical, 17:596–606.

Bassi, L., Ricci, D., Volzone, A., Allsop, J. M., Srinivasan, L., Pai, A., Ribes, C., Ramenghi, L. A., Mercuri, E., Mosca, F., et al. (2008). Probabilistic diffusion tractography of the optic radiations and visual function in preterm infants at term equivalent age. Brain, 131(2):573–582.

Batalle, D., Hughes, E. J., Zhang, H., Tournier, J.-D., Tusor, N., Aljabar, P., Wali, L., Alexander, D. C., Hajnal, J. V., Nosarti, C., et al. (2017). Early development of structural networks and the impact of prematurity on brain connectivity. Neuroimage, 149:379–392.

Batalle, D., O’Muircheartaigh, J., Makropoulos, A., Kelly, C. J., Dimitrova, R., Hughes, E. J., Hajnal, J. V., Zhang, H., Alexander, D. C., Edwards, A. D., et al. (2019). Different patterns of cortical maturation before and after 38 weeks gestational age demonstrated by diffusion mri in vivo. NeuroImage, 185:764–775.

Baykara, E., Gesierich, B., Adam, R., Tuladhar, A. M., Biesbroek, J. M., Koek, H. L., Ropele, S., Jouvent, E., Initiative, A. D. N., Chabriat, H., et al. (2016). A novel imaging marker for small vessel disease based on skeletonization of white matter tracts and diffusion histograms. Annals of neurology, 80(4):581–592.

Beck, S., Wojdyla, D., Say, L., Betran, A. P., Merialdi, M., Requejo, J. H., Rubens, C., Menon, R., and Van Look, P. F. (2010). The worldwide incidence of preterm birth: a systematic review of maternal mortality and morbidity. Bulletin of the World Health Organization, 88:31–38.

Benjamini, Y. and Hochberg, Y. (1995). Controlling the false discovery rate: a practical and powerful approach to multiple testing. Journal of the Royal statistical society: series B (Methodological), 57(1):289–300.

Bhutta, A. T., Cleves, M. A., Casey, P. H., Cradock, M. M., and Anand, K. J. (2002). Cognitive and behavioral outcomes of school-aged children who were born preterm: a meta-analysis. Jama, 288(6):728–737.

Bilenko, N. Y. and Gallant, J. L. (2016). Pyrcca: regularized kernel canonical correlation analysis in python and its applications to neuroimaging. Frontiers in neuroinformatics, 10:49.

Blencowe, H., Cousens, S., Chou, D., Oestergaard, M., Say, L., Moller, A.-B., Kinney, M., and Lawn, J. (2013). Born too soon: the global epidemiology of 15 million preterm births. Reproductive health, 10(1):1–14.

Blesa, M., Galdi, P., Sullivan, G., Wheater, E. N., Stoye, D. Q., Lamb, G. J., Quigley, A. J., Thrippleton, M. J., Bastin, M. E., and Boardman, J. P. (2020). Peak width of skeletonized water diffusion mri in the neonatal brain. Frontiers in Neurology, 11:235.

Chamberland, M., Raven, E. P., Genc, S., Duffy, K., Descoteaux, M., Parker, G. D., Tax, C. M., and Jones, D. K. (2019). Dimensionality reduction of diffusion mri measures for improved tractometry of the human brain. NeuroImage, 200:89–100.

Chin, R., You, A. X., Meng, F., Zhou, J., and Sim, K. (2018). Recognition of schizophrenia with regularized support vector machine and sequential region of interest selection using structural magnetic resonance imaging. Scientific Reports, 8(1):1–10.

Chu, C., Lagercrantz, H., Forssberg, H., and Nagy, Z. (2015). Investigating the use of support vector machine classification on structural brain images of preterm–born teenagers as a biological marker. Plos one, 10(4):e0123108.

Collins, S. E., Spencer-Smith, M., Mürner-Lavanchy, I., Kelly, C. E., Pyman, P., Pascoe, L., Cheong, J., Doyle, L. W., Thompson, D. K., and Anderson, P. J. (2019). White matter microstructure correlates with mathematics but not word reading performance in 13-year-old children born very preterm and full-term. NeuroImage: Clinical, 24:101944.

Correa, N. M., Adali, T., Li, Y.-O., and Calhoun, V. D. (2010a). Canonical correlation analysis for data fusion and group inferences. IEEE signal processing magazine, 27(4):39–50.

Correa, N. M., Eichele, T., Adali, T., Li, Y.-O., and Calhoun, V. D. (2010b). Multi-set canonical correlation analysis for the fusion of concurrent single trial erp and functional mri. Neuroimage, 50(4):1438–1445.

Correa, N. M., Li, Y.-O., Adali, T., and Calhoun, V. D. (2008). Canonical correlation analysis for feature-based fusion of biomedical imaging modalities and its application to detection of associative networks in schizophrenia. IEEE journal of selected topics in signal processing, 2(6):998–1007.

Counsell, S. J., Allsop, J. M., Harrison, M. C., Larkman, D. J., Kennea, N. L., Kapellou, O., Cowan, F. M., Hajnal, J. V., Edwards, A. D., and Rutherford, M. A. (2003). Diffusion-weighted imaging of the brain in preterm infants with focal and diffuse white matter abnormality. Pediatrics, 112(1):1–7.

Counsell, S. J., Edwards, A. D., Chew, A. T., Anjari, M., Dyet, L. E., Srinivasan, L., Boardman, J. P., Allsop, J. M., Hajnal, J. V., Rutherford, M. A., et al. (2008). Specific relations between neurodevelopmental abilities and white matter microstructure in children born preterm. Brain, 131(12):3201–3208.

Counsell, S. J., Maalouf, E. F., Fletcher, A. M., Duggan, P., Battin, M., Lewis, H. J., Herlihy, A. H., Edwards, A. D., Bydder, G. M., and Rutherford, M. A. (2002). Mr imaging assessment of myelination in the very preterm brain. American journal of neuroradiology, 23(5):872–881.

Cox, S. R., Ritchie, S. J., Tucker-Drob, E. M., Liewald, D. C., Hagenaars, S. P., Davies, G., Wardlaw, J. M., Gale, C. R., Bastin, M. E., and Deary, I. J. (2016). Ageing and brain white matter structure in 3,513 uk biobank participants. Nature communications, 7(1):1–13.

Daducci, A., Canales-Rodríguez, E. J., Zhang, H., Dyrby, T. B., Alexander, D. C., and Thiran, J.-P. (2015). Accelerated microstructure imaging via convex optimization (amico) from diffusion mri data. Neuroimage, 105:32–44.

Davatzikos, C. (2019). Machine learning in neuroimaging: Progress and challenges. Neuroimage, 197:652.

De Santis, S., Drakesmith, M., Bells, S., Assaf, Y., and Jones, D. K. (2014). Why diffusion tensor mri does well only some of the time: variance and covariance of white matter tissue microstructure attributes in the living human brain. Neuroimage, 89:35–44.

Dhollander, T., Mito, R., Raffelt, D., and Connelly, A. (2019). Improved white matter response function estimation for 3-tissue constrained spherical deconvolution. In Proc. Intl. Soc. Mag. Reson. Med, volume 555.

Dhollander, T., Raffelt, D., and Connelly, A. (2016). Unsupervised 3-tissue response function estimation from single-shell or multi-shell diffusion mr data without a co-registered t1 image. In ISMRM Workshop on Breaking the Barriers of Diffusion MRI, volume 5. ISMRM.

Dhollander, T., Zanin, J., Nayagam, B. A., Rance, G., and Connelly, A. (2018). Feasibility and benefits of 3-tissue constrained spherical deconvolution for studying the brains of babies. In Proceedings of the 26th annual meeting of the International Society of Magnetic Resonance in Medicine, page 3077.

Doshi, J., Erus, G., Ou, Y., Gaonkar, B., and Davatzikos, C. (2013). Multi-atlas skull-stripping. Academic radiology, 20(12):1566–1576.

Duerden, E., Foong, J., Chau, V., Branson, H., Poskitt, K., Grunau, R., Synnes, A., Zwicker, J., and Miller, S. (2015). Tract-based spatial statistics in preterm-born neonates predicts cognitive and motor outcomes at 18 months. American Journal of Neuroradiology, 36(8):1565–1571.

Dyet, L. E., Kennea, N., Counsell, S. J., Maalouf, E. F., Ajayi-Obe, M., Duggan, P. J., Harrison, M., Allsop, J. M., Hajnal, J., Herlihy, A. H., et al. (2006). Natural history of brain lesions in extremely preterm infants studied with serial magnetic resonance imaging from birth and neurodevelopmental assessment. Pediatrics, 118(2):536–548.

Eaton-Rosen, Z., Melbourne, A., Orasanu, E., Cardoso, M. J., Modat, M., Bainbridge, A., Kendall, G. S., Robertson, N. J., Marlow, N., and Ourselin, S. (2015). Longitudinal measurement of the developing grey matter in preterm subjects using multi-modal mri. NeuroImage, 111:580–589.

Fadnavis, S., Batson, J., and Garyfallidis, E. (2020). Patch2self: Denoising diffusion mri with self-supervised learning. Advances in Neural Information Processing Systems, 33:16293–16303.

Galdi, P., Blesa, M., Stoye, D. Q., Sullivan, G., Lamb, G. J., Quigley, A. J., Thrippleton, M. J., Bastin, M. E., and Boardman, J. P. (2020). Neonatal morphometric similarity mapping for predicting brain age and characterizing neuroanatomic variation associated with preterm birth. NeuroImage: Clinical, 25:102195.

Gaonkar, B. and Davatzikos, C. (2013). Analytic estimation of statistical significance maps for support vector machine based multi-variate image analysis and classification. Neuroimage, 78:270–283.

Garyfallidis, E., Brett, M., Amirbekian, B., Rokem, A., Van Der Walt, S., Descoteaux, M., and Nimmo-Smith, I. (2014). Dipy, a library for the analysis of diffusion mri data. Frontiers in neuroinformatics, 8:8.

Girault, J. B., Munsell, B. C., Puechmaille, D., Goldman, B. D., Prieto, J. C., Styner, M., and Gilmore, J. H. (2019). White matter connectomes at birth accurately predict cognitive abilities at age 2. Neuroimage, 192:145–155.

Golland, P., Fischl, B., Spiridon, M., Kanwisher, N., Buckner, R. L., Shenton, M. E., Kikinis, R., Dale, A., and Grimson, W. E. L. (2002). Discriminative analysis for image-based studies. In International Conference on Medical Image Computing and Computer-Assisted Intervention, pages 508–515. Springer.

Groppo, M., Ricci, D., Bassi, L., Merchant, N., Doria, V., Arichi, T., Allsop, J. M., Ramenghi, L., Fox, M. J., Cowan, F. M., et al. (2014). Development of the optic radiations and visual function after premature birth. Cortex, 56:30–37.

Hardoon, D. R., Mourao-Miranda, J., Brammer, M., and Shawe-Taylor, J. (2007). Unsupervised analysis of fmri data using kernel canonical correlation. NeuroImage, 37(4):1250–1259.

Hardoon, D. R., Szedmak, S., and Shawe-Taylor, J. (2004). Canonical correlation analysis: An overview with application to learning methods. Neural computation, 16(12):2639–2664.

Hosseini, F., Ebrahimpourkomleh, H., and KhodamHazrati, M. (2015). Quantitative evaluation of skull stripping techniques on magnetic resonance images. In World Congress on Electrical Engineering and Computer Systems and Science (EECSS 2015).

Hughes, E., Cordero-Grande, L., Murgasova, M., Hutter, J., Price, A., Gomes, A. D. S., Allsop, J., Steinweg, J., Tusor, N., Wurie, J., et al. (2017). The developing human connectome: announcing the first release of open access neonatal brain imaging. Organization for Human Brain Mapp, pages 25–29.

Hüppi, P S., Maier, S. E., Peled, S., Zientara, G. P., Barnes, P. D., Jolesz, F. A., and Volpe, J. J. (1998a). Microstructural development of human newborn cerebral white matter assessed in vivo by diffusion tensor magnetic resonance imaging. Pediatric research, 44(4):584–590.

Hüppi, P. S., Warfield, S., Kikinis, R., Barnes, P. D., Zientara, G. P., Jolesz, F. A., Tsuji, M. K., and Volpe, J. J. (1998b). Quantitative magnetic resonance imaging of brain development in premature and mature newborns. Annals of Neurology: Official Journal of the American Neurological Association and the Child Neurology Society, 43(2):224–235.

Iglesias, J. E., Liu, C.-Y., Thompson, P. M., and Tu, Z. (2011). Robust brain extraction across datasets and comparison with publicly available methods. IEEE transactions on medical imaging, 30(9):1617–1634.

Jensen, J. H., Helpern, J. A., Ramani, A., Lu, H., and Kaczynski, K. (2005). Diffusional kurtosis imaging: the quantification of non-gaussian water diffusion by means of magnetic resonance imaging. Magnetic Resonance in Medicine: An Official Journal of the International Society for Magnetic Resonance in Medicine, 53(6):1432–1440.

Jeurissen, B., Tournier, J.-D., Dhollander, T., Connelly, A., and Sijbers, J. (2014). Multi-tissue constrained spherical deconvolution for improved analysis of multi-shell diffusion mri data. NeuroImage, 103:411–426.

Jones, D. K. and Cercignani, M. (2010). Twenty-five pitfalls in the analysis of diffusion mri data. NMR in Biomedicine, 23(7):803–820.

Kaden, E., Kruggel, F., and Alexander, D. C. (2016). Quantitative mapping of the per-axon diffusion coefficients in brain white matter. Magnetic resonance in medicine, 75(4):1752–1763.

Kelly, C. E., Thompson, D. K., Chen, J., Leemans, A., Adamson, C. L., Inder, T. E., Cheong, J. L., Doyle, L. W., and Anderson, P. J. (2016). Axon density and axon orientation dispersion in children born preterm. Human brain mapping, 37(9):3080–3102.

Kimpton, J., Batalle, D., Barnett, M., Hughes, E., Chew, A., Falconer, S., Tournier, J., Alexander, D., Zhang, H., Edwards, A., et al. (2021). Diffusion magnetic resonance imaging assessment of regional white matter maturation in preterm neonates. Neuroradiology, 63(4):573–583.

Knight, M. J., Smith-Collins, A., Newell, S., Denbow, M., and Kauppinen, R. A. (2018). Cerebral white matter maturation patterns in preterm infants: an mri t2 relaxation anisotropy and diffusion tensor imaging study. Journal of Neuroimaging, 28(1):86–94.

Kulikova, S., Hertz-Pannier, L., Dehaene-Lambertz, G., Buzmakov, A., Poupon, C., and Dubois, J. (2015). Multi-parametric evaluation of the white matter maturation. Brain Structure and Function, 220(6):3657–3672.

Kunz, N., Zhang, H., Vasung, L., O’Brien, K. R., Assaf, Y., Lazeyras, F., Alexander, D. C., and Hüppi, P. S. (2014). Assessing white matter microstructure of the newborn with multi-shell diffusion mri and biophysical compartment models. Neuroimage, 96:288–299.

Lao, Z., Shen, D., Xue, Z., Karacali, B., Resnick, S. M., and Davatzikos, C. (2004). Morphological classification of brains via high-dimensional shape transformations and machine learning methods. Neuroimage, 21(1):46–57.

Ling, X., Tang, W., Liu, G., Huang, L., Li, B., Li, X., Liu, S., and Xu, J. (2013). Assessment of brain maturation in the preterm infants using diffusion tensor imaging (dti) and enhanced t2 star weighted angiography (eswan). European Journal of Radiology, 82(9):e476–e483.

Ly, M. T., Nanavati, T. U., Frum, C. A., and Pergami, P. (2015). Comparing tract-based spatial statistics and manual region-of-interest labeling as diffusion analysis methods to detect white matter abnormalities in infants with hypoxic-ischemic encephalopathy. Journal of Magnetic Resonance Imaging, 42(6):1689–1697.

Mürner-Lavanchy, I. M., Kelly, C. E., Reidy, N., Doyle, L. W., Lee, K. J., Inder, T., Thompson, D. K., Morgan, A. T., and Anderson, P. J. (2018). White matter microstructure is associated with language in children born very preterm. NeuroImage: Clinical, 20:808–822.

Ouyang, M., Dubois, J., Yu, Q., Mukherjee, P., and Huang, H. (2019a). Delineation of early brain development from fetuses to infants with diffusion mri and beyond. Neuroimage, 185:836–850.

Ouyang, M., Jeon, T., Sotiras, A., Peng, Q., Mishra, V., Halovanic, C., Chen, M., Chalak, L., Rollins, N., Roberts, T. P., et al. (2019b). Differential cortical microstructural maturation in the preterm human brain with diffusion kurtosis and tensor imaging. Proceedings of the National Academy of Sciences, 116(10):4681–4688.

Pannek, K., Fripp, J., George, J. M., Fiori, S., Colditz, P. B., Boyd, R. N., and Rose, S. E. (2018). Fixel-based analysis reveals alterations is brain microstructure and macrostructure of preterm-born infants at term equivalent age. NeuroImage: Clinical, 18:51–59.

Partridge, S. C., Mukherjee, P., Henry, R. G., Miller, S. P., Berman, J. I., Jin, H., Lu, Y., Glenn, O. A., Ferriero, D. M., Barkovich, A. J., et al. (2004). Diffusion tensor imaging: serial quantitation of white matter tract maturity in premature newborns. Neuroimage, 22(3):1302–1314.

Pecheva, D., Kelly, C., Kimpton, J., Bonthrone, A., Batalle, D., Zhang, H., and Counsell, S. (2018). Recent advances in diffusion neuroimaging: applications in the developing preterm brain. f1000res. F1000FacultyRev-1326.

Pecheva, D., Yushkevich, P., Batalle, D., Hughes, E., Aljabar, P., Wurie, J., Hajnal, J. V., Edwards, A. D., Alexander, D. C., Counsell, S. J., et al. (2017). A tract-specific approach to assessing white matter in preterm infants. Neuroimage, 157:675–694.

Saha, S., Pagnozzi, A., Bourgeat, P., George, J. M., Bradford, D., Colditz, P. B., Boyd, R. N., Rose, S. E., Fripp, J., and Pannek, K. (2020). Predicting motor outcome in preterm infants from very early brain diffusion mri using a deep learning convolutional neural network (cnn) model. Neuroimage, 215:116807.

Schilling, K. G., Fadnavis, S., Batson, J., Visagie, M., Combes, A. J., By, S., McKnight, C. D., Bagnato, F., Garyfallidis, E., Landman, B. A., et al. (2022). Denoising of diffusion mri in the cervical spinal cord-effects of denoising strategy and acquisition on intra-cord contrast, signal modeling, and feature conspicuity. NeuroImage, page 119826.

Schölkopf, B., Smola, A. J., Bach, F., et al. (2002). Learning with kernels: support vector machines, regularization, optimization, and beyond. MIT press.

Shattuck, D. W. and Leahy, R. M. (2000). Brainsuite: An automated cortical surface identification tool. In International Conference on Medical Image Computing and Computer-Assisted Intervention, pages 50–61. Springer.

Shi, J., Chang, L., Wang, J., Zhang, S., Yao, Y., Zhang, S., Jiang, R., Guo, L., Guan, H., and Zhu, W. (2016). Initial application of diffusional kurtosis imaging in evaluating brain development of healthy preterm infants. PLoS One, 11(4):e0154146.

Smith, S. M. (2000). Bet: Brain extraction tool. FMRIB TR00SMS2b, Oxford Centre for Functional Magnetic Resonance Imaging of the Brain), Department of Clinical Neurology, Oxford University, John Radcliffe Hospital, Headington, UK.

Smith, S. M., Jenkinson, M., Johansen-Berg, H., Rueckert, D., Nichols, T. E., Mackay, C. E., Watkins, K. E., Ciccarelli, O., Cader, M. Z., Matthews, P. M., et al. (2006). Tract-based spatial statistics: voxelwise analysis of multi-subject diffusion data. Neuroimage, 31(4):1487–1505.

Sui, J., He, H., Yu, Q., Chen, J., Rogers, J., Pearlson, G. D., Mayer, A., Bustillo, J., Canive, J., and Calhoun, V. D. (2013). Combination of resting state fmri, dti, and smri data to discriminate schizophrenia by n-way mcca+ jica. Frontiers in human neuroscience, 7:235.

Sui, J., Pearlson, G., Caprihan, A., Adali, T., Kiehl, K. A., Liu, J., Yamamoto, J., and Calhoun, V. D. (2011). Discriminating schizophrenia and bipolar disorder by fusing fmri and dti in a multimodal cca+ joint ica model. Neuroimage, 57(3):839–855.

Thompson, D. K., Inder, T. E., Faggian, N., Johnston, L., Warfield, S. K., Anderson, P. J., Doyle, L. W., and Egan, G. F. (2011). Characterization of the corpus callosum in very preterm and full-term infants utilizing mri. Neuroimage, 55(2):479–490.

Thompson, D. K., Kelly, C. E., Chen, J., Beare, R., Alexander, B., Seal, M. L., Lee, K., Matthews, L. G., Anderson, P. J., Doyle, L. W., et al. (2019a). Early life predictors of brain development at term-equivalent age in infants born across the gestational age spectrum. Neuroimage, 185:813–824.

Thompson, D. K., Kelly, C. E., Chen, J., Beare, R., Alexander, B., Seal, M. L., Lee, K. J., Matthews, L. G., Anderson, P. J., Doyle, L. W., et al. (2019b). Characterisation of brain volume and microstructure at term-equivalent age in infants born across the gestational age spectrum. NeuroImage: Clinical, 21:101630.

Timmers, I., Roebroeck, A., Bastiani, M., Jansma, B., Rubio-Gozalbo, E., and Zhang, H. (2016). Assessing microstructural substrates of white matter abnormalities: a comparative study using dti and noddi. PloS one, 11(12):e0167884.

Tokariev, M., Vuontela, V., Perkola, J., Lönnberg, P., Lano, A., Andersson, S., Metsäranta, M., and Carlson, S. (2020). A protocol for the analysis of dti data collected from young children. MethodsX, 7:100878.

Tortora, D., Martinetti, C., Severino, M., Uccella, S., Malova, M., Parodi, A., Brera, F., Morana, G., Ramenghi, L. A., and Rossi, A. (2018). The effects of mild germinal matrix-intraventricular haemorrhage on the developmental white matter microstructure of preterm neonates: a dti study. European radiology, 28(3):1157–1166.

Tournier, J.-D., Smith, R., Raffelt, D., Tabbara, R., Dhollander, T., Pietsch, M., Christiaens, D., Jeurissen, B., Yeh, C.-H., and Connelly, A. (2019). Mrtrix3: A fast, flexible and open software framework for medical image processing and visualisation. Neuroimage, 202:116137.

Tustison, N. J., Avants, B. B., Cook, P. A., Zheng, Y., Egan, A., Yushkevich, P. A., and Gee, J. C. (2010). N4itk: improved n3 bias correction. IEEE transactions on medical imaging, 29(6):1310–1320.

Vaher, K., Galdi, P., Cabez, M. B., Sullivan, G., Stoye, D. Q., Quigley, A. J., Thrippleton, M. J., Bogaert, D., Bastin, M. E., Cox, S. R., et al. (2022). General factors of white matter microstructure from dti and noddi in the developing brain. NeuroImage, 254:119169.

Vapnik, V. N. (1999). An overview of statistical learning theory. IEEE transactions on neural networks, 10(5):988–999.

Volpe, J. J. (2003). Cerebral white matter injury of the premature infant—more common than you think. Pediatrics, 112(1):176–180.

Wang, H.-T., Smallwood, J., Mourao-Miranda, J., Xia, C. H., Satterthwaite, T. D., Bassett, D. S., and Bzdok, D. (2020). Finding the needle in a high-dimensional haystack: Canonical correlation analysis for neuroscientists. NeuroImage, 216:116745.

Whitwell, J. L. (2009). Voxel-based morphometry: an automated technique for assessing structural changes in the brain. Journal of Neuroscience, 29(31):9661–9664.

Young, J. M., Vandewouw, M. M., Mossad, S. I., Morgan, B. R., Lee, W., Smith, M. L., Sled, J. G., and Taylor, M. J. (2019). White matter microstructural differences identified using multi-shell diffusion imaging in six-year-old children born very preterm. NeuroImage: Clinical, 23:101855.

Zhang, H., Schneider, T., Wheeler-Kingshott, C. A., and Alexander, D. C. (2012). Noddi: practical in vivo neurite orientation dispersion and density imaging of the human brain. Neuroimage, 61(4):1000–1016.

Zhao, X., Zhang, C., Zhang, B., Yan, J., Wang, K., Zhu, Z., and Zhang, X. (2021). The value of diffusion kurtosis imaging in detecting delayed brain development of premature infants.

