## Supplementary Figures for "Data-driven characterization of Preterm Birth through intramodal Diffusion MRI"

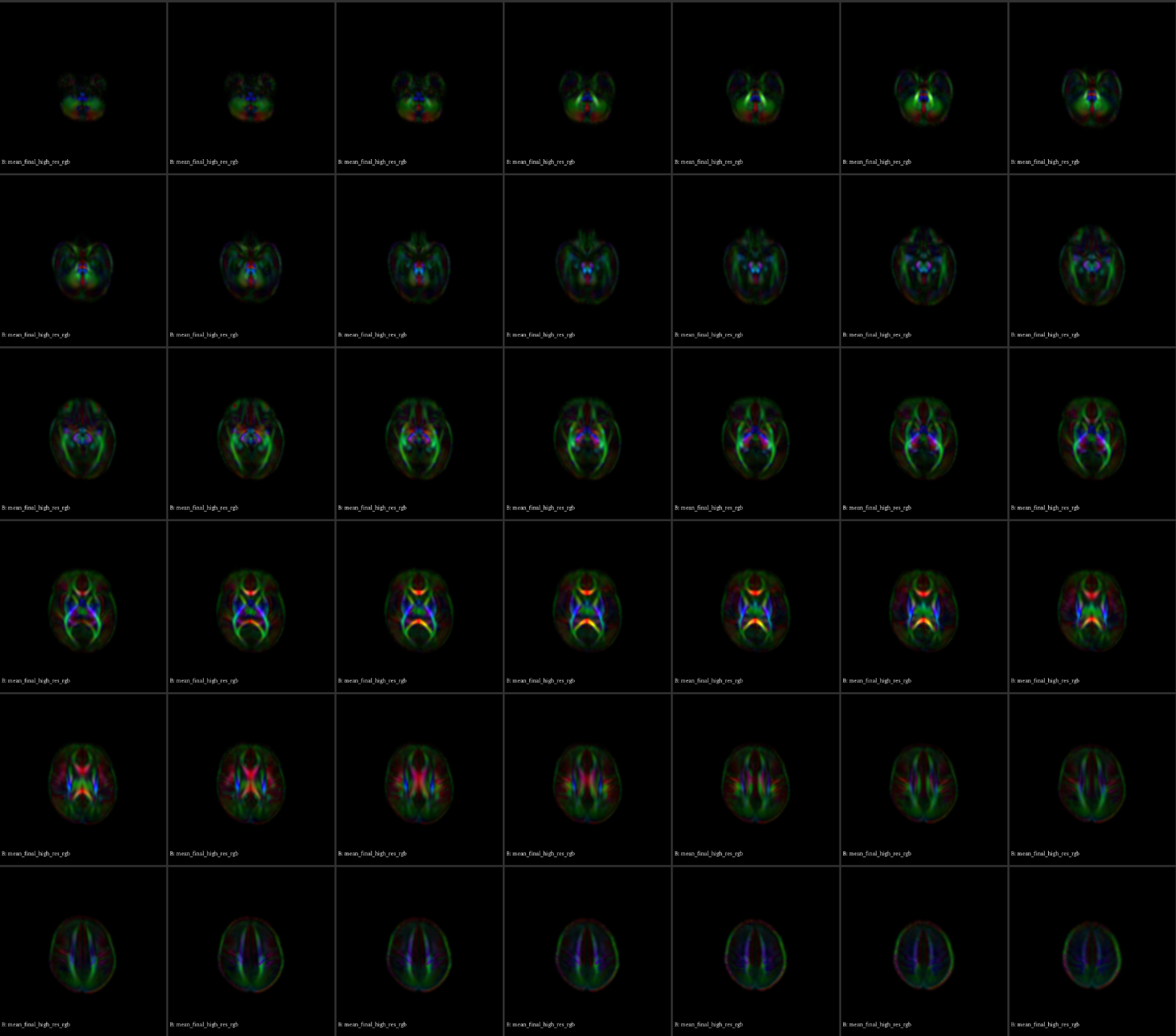

**Figure S1.** Population-specific DTI template: lightbox displaying axial views of the age-specific template created ad-hoc for performing normalization of DTI volumes within DTI-TK and subsequent WM skeleton creation within TBSS.

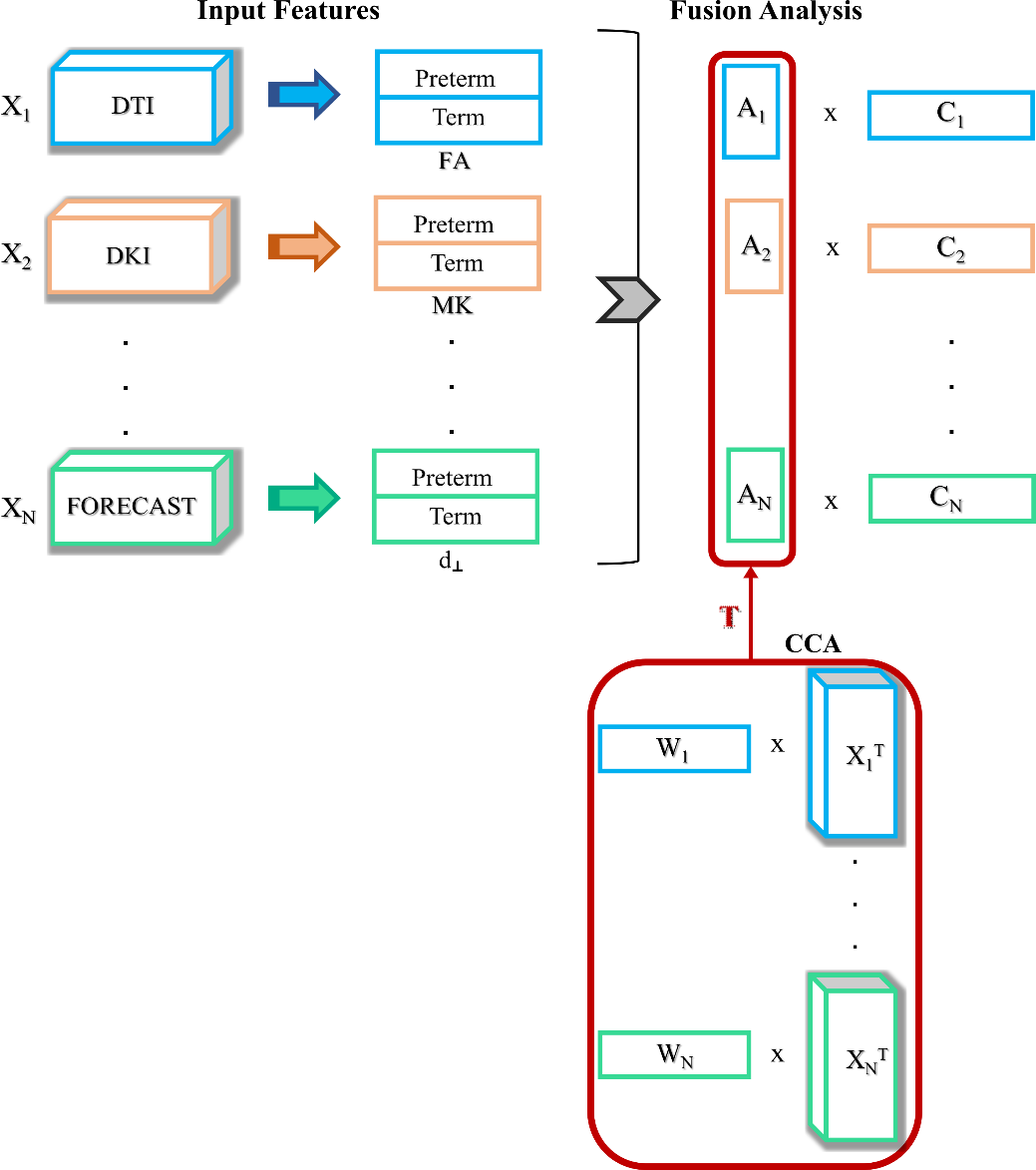

**Figure S2.** Canonical Correlation Analysis framework applied to our intramodal dataset on the all the 14 HARDI microstructural measures
